# Lipid gating of BK channels and mechanism of activation by negatively charged lipids

**DOI:** 10.1101/2025.05.27.656279

**Authors:** Andrei Mironenko, Bert L. de Groot, Wojciech Kopec

## Abstract

BK channels are a class of K^+^ channels that possess an unusually high conductance and are synergistically gated by intracellular Ca^2+^ and voltage. Despite the significant array of experimental and computational data, many aspects of their function and dynamics remain unclear - such as how ion permeation is halted in the closed state of the channel. Available CryoEM structures obtained in deactivating conditions capture the channel with a wide, unobstructed pore, in contrast to e.g. a helix bundle crossing observed in some K^+^ channels. Several hypotheses of BK closure were proposed, including selectivity filter and hydrophobic gating. In this work, we expand on the model of hydrophobic gating and focus on the role of lipids. Using atomistic molecular dynamics simulations with applied voltage, we directly investigate the ability of the full length BK channel in various CryoEM states to permeate ions, and propose lipid entrance into the pore - either with lipid tails or entire lipid molecules - through the membrane-facing fenestrations to be a critical determinant of BK conductivity. Furthermore, we elucidate the mechanism of BK activation by negatively charged lipids via a combination of atomistic & coarse-grained simulations, and suggest that they act by a multi-modal mechanism, which encompasses lipid entry reduction, increase of the K^+^ occupancy of the pore, and stabilization of the channel’s open-state structure - in broad agreement with experimental data.

## Introduction

Potassium (K^+^) channels enable permeation of K^+^ ions across biological membranes with high conductance and high selectivity, which plays an important role in a variety of cellular processes ^1–3^. In particular, BK channels (‘big K^+^’, also known as maxiK or Slo1) stand out among other K^+^ channels due to their particularly large single-channel conductance - 100-300 pS ^4^. BK channels are activated synergistically by intracellular Ca^2+^ and membrane depolarization; this allows them to participate in a plethora of important physiological functions across various tissues, e.g. maintenance of smooth muscle arterial tone, K^+^ secretion in kidneys and termination of action potential in neurons ^5–7^.

The first full-length CryoEM structures were obtained for the BK channel from *Aplysia californica* ^8,9^, followed by structures from other organisms (including human); some of the structures also featured additional regulatory subunits ^10–12^. The structures revealed that the pore-forming ɑ-subunits of BK channels, in addition to a pore domain, contain a transmembrane voltage sensing domain, as well as an intracellular Ca^2+^-sensing domain that together enable the synergistic activation (*Fig. 1A*). In certain cases, CryoEM structures were obtained either in presence, or absence of Ca^2+^, allowing to isolate the putatively open and closed states of the channel, respectively ^8–11^. However, whether these structures actually correspond to open (conductive) or closed (non-conductive) states has been unclear, especially considering the fact that neither structure had a physical constriction of the permeation pathway that would prevent ion permeation in the closed state ^8,9,11^ - contrary to e.g. Kv channels, where a helix-bundle crossing sterically blocks the pore ^13,14^. In a recent CryoEM study, mutations of voltage-sensing arginines that promote deactivation made it possible to obtain structures with the pore as narrow as ∼8 Å in diameter ^12^, which is nevertheless wide enough that it’s sterically possible to fit a hydrated K^+^ ion ^15^.

**Fig. 1.**
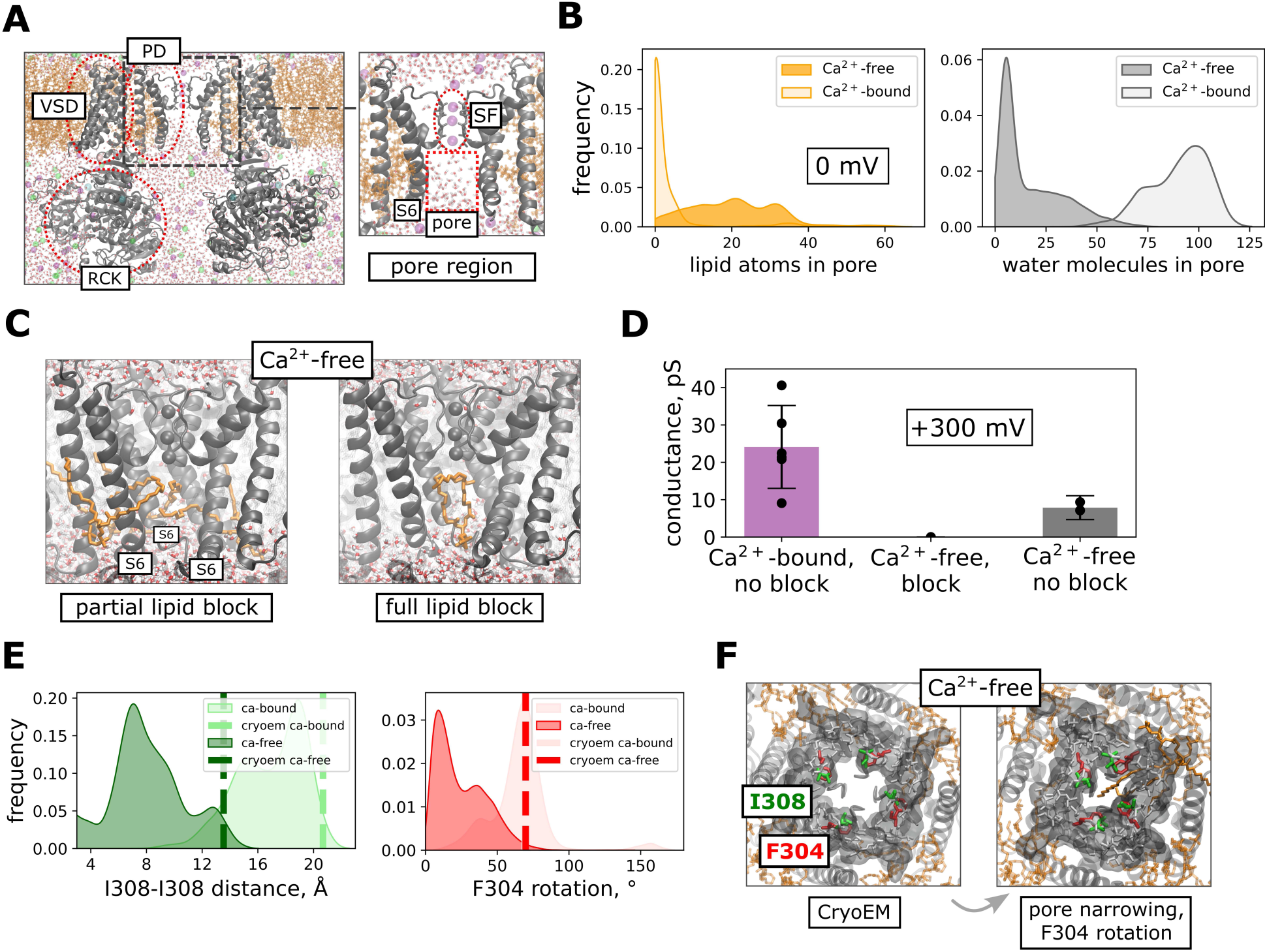
(*A*) BK channels feature the pore domain (PD), the voltage-sensing domain (VSD), and the cytosolic Ca^2+^-sensing domain (RCK) (left); a zoom-in on the pore region with the pore cavity, lined by S6 helices with the selectivity filter above (right). A snapshot from one of our atomistic simulations of BK, with the channel in the lipid bilayer, water and ions included in simulations is shown; only two opposing domains out of four are shown for clarity (*B*) Distribution of the number of lipid atoms (non-hydrogen atoms out of ∼50 per lipid molecule) and water molecules in the pore over all replicas in the all-atom simulations of Ca^2+^-free and Ca^2+^-bound without applied voltage. (*C*) Snapshots showing the two types of lipid block we observed in our simulations of the Ca^2+^-free state: partial (only lipid tails are in the pore) and full lipid block (entire lipid molecules occupy the pore). For clarity, one out of four subunits, which would be located in the front, is not shown. (*D*) Conductances of Ca^2+^-free and Ca^2+^-bound under applied voltage (+300 mV). ‘Ca^2+^-free, block’ includes all applied voltage simulations started with either partial or full lipid block; additionally, conductance in simulations of Ca^2+^-free where no lipid block occurred is shown. (*E*) Distributions of pore widths, defined as distance between I308 residues of opposite subunits, and rotation of F304 into the pore as the angle between the center of mass of F304 from all subunits, and CA and CZ of a F304 from each subunit. Dashed lines represent values in respective CryoEM structures; F304 angle values are similar in both CryoEM structures. (*F*) Snapshots from simulations of Ca^2+^-free showing the conformational changes relative to the CryoEM structures, including narrowing of the pore at I308 and inward rotation of F304 into the pore, necessary to achieve the non-conductive state of the channel.

While it is possible that BK conformations with a fully constricted pore might exist, albeit not captured by the currently available structures - mechanisms other than physical block by helix-bundle-like structures have also been proposed. One such possibility is gating at the selectivity filter (SF). The SF is the narrowest part of the ion permeation pathway in K^+^ channels that largely determines the channel’s conductance and selectivity ^16^. In some K^+^ channels, ion permeation can be stopped via conformational changes at the SF, as is the case in several K2P channels ^17,18^. While in BK the SF is in the same (conductive) conformation in both Ca^2+^-bound and Ca^2+^-free CryoEM structures ^8,9^, it was shown that large organic molecules are able to block the pore not only in the open, but in the closed state as well, raising the possibility that the channel is gated at the SF ^19–23;^ to note, some data support such block only in the partially closed state, but not fully closed ^19^. In another study, molecular dynamics (MD) simulations of the human BK channel showed destabilization of the SF in the Ca^2+^-free-like state ^24^, although it should be noted that a homology model was used for the structure of the channel, which, together with an initial configuration containing water molecules in the SF, could promote destabilization of the SF structure ^16^.

An alternative hypothesis is hydrophobic gating. In this mechanism, water molecules leave the inner pore of the channel in the closed state, creating a hydrophobic barrier for ion permeation through the pore ^25^. Hydrophobic gating was first proposed for model nanopores based on MD simulations ^26,27^: even if the diameter of a narrow pore was bigger than that of a water molecule, such pores could still undergo spontaneous dehydration-hydration transitions. There’s structural and computational evidence of hydrophobic gating in several biological ion channels as well (e.g. GLIC ^28,29^, MscS ^30–32^, or TWIK-1 ^33^). For BK, several MD studies showed cavity dehydration in the Ca^2+^-free state, and a hydrated pore in the Ca^2+^-bound state ^34–36;^ additionally, the Ca^2+^-free structure underwent further pore narrowing and rotation of pore-lining residues, proposed to increase the hydrophobicity of the pore ^34^. The idea of pore dehydration in BK agrees well with the experimental mutagenesis data, where the voltage of half activation correlates with the hydrophobicity of pores of various mutants ^35–39^.

Despite the overall support of the hydrophobic gating model in BK channels by experimental and computational data, a reoccurring topic in studies is regulation of hydrophobic gating of BK by lipids. In several MD studies of BK channels pore dehydration was accompanied by entrance of lipid tails into the pore through the fenestrations between the pore-forming S6 helices ^12,34–36^, further addressed in a recent computational study that showed facilitation of pore dehydration by penetrating lipids ^40^. This is especially interesting, as the mentioned fenestrations occur only in the CryoEM structures obtained in deactivating conditions, but not in activating ^8,9,11^, with some CryoEM structures also capturing lipid tails in these fenestrations ^11^. At the same time, the inaccessibility of the BK pore to blockers in the fully closed state observed in an experimental study ^19^ could indicate the existence of a steric block of the pore, which could be posed either by the channel itself, or by lipids occupying the pore. Additionally, the membrane lipid composition has a strong effect on BK channel’s activity - it was shown to be regulated by the lipid’s tail length ^41^, as well as head group charge and size ^42,43^, prompting further investigation into the mechanistic underpinnings of this effect, as well as a potential role of lipid composition in the suggested lipid dynamics in the pore.

In this work, we focus on the role of lipids in BK gating using atomistic and coarse-grained MD simulations. To our knowledge, the ion permeability of either a hydrated or dehydrated pore has been tested directly using MD simulations only for the isolated pore domain of BK, which was shown to be conformationally unstable ^35^ - but not in the full-length channel. Thus, we first test ion permeabilities of the full-length BK CryoEM structures using simulations with applied voltage. We describe the various modes of lipid behavior we observed: in particular, partial and full entry of lipids into the pore, as well as the interplay between lipid entry and conformational changes in the protein and its effect on the channel’s conductance. Finally, we investigate BK activation by negatively charged lipids, and suggest a multi-modal mechanism which involves effects on the channel’s conformation as well as lipid dynamics in the pore.

## Results

### All-atom simulations of Ca^2+^-bound and Ca^2+^-free states

Previously, it was shown that Ca^2+^-free states of BK generally undergo cavity dehydration in MD simulations, and the pore in Ca^2+^-bound channels remains hydrated ^34–36;^ additionally, a recent MD study used applied voltage to test the conductivity of the isolated pore domain of BK, although without special restraints to prevent lipids from entering the pore the channel’s tetrameric structure collapsed ^35^. Therefore, the ability of the CryoEM states of the full-length channel to conduct ions has not been confirmed directly. To bridge this gap, we first performed atomistic MD simulations of BK from *Aplysia californica* in equilibrium (i.e. at 0 mV), and then under applied voltage starting from various frames of the equilibrium runs, in both Ca^2+^-bound and Ca^2+^-free states (PDB ID 5tj6 ^9^ and 5tji ^8^, respectively; +300 mV, POPC membrane, 0.8 M KCl, CHARMM36m). The simulation durations for each condition tested are given in *SI Table 1*.

The pore of Ca^2+^-bound stayed hydrated throughout most of the equilibrium and applied voltage simulations in agreement with previous studies (*Fig. 1B*), with the exception of a decrease in pore hydration in one equilibrium replica (an independent repeat of simulations) out of 5 (*SI Fig. 1B*), and spontaneous dehydration events under applied voltage in 2 out of 6 replicas (*SI Fig. 3*). Conversely, the Ca^2+^-free channel showed a mostly dehydrated pore (*Fig. 1B*), with the exception of one equilibrium replica (out of 5), and all the applied voltage simulations started from this replica (*SI Fig. 1A, C*). Notably, pore dehydration was always accompanied by lipids entering into the pore. We observed two types of such events: in the first type, only lipid tails entered through the membrane-exposed fenestrations between S6 helices of neighboring subunits present in the Ca^2+^-free structure (‘partial’ lipid block, *Fig. 1C*), as in the case of several replicas of the Ca^2+^-free state (*, SI Fig. 1A, D*). The second type was the entrance of entire lipid molecules through the fenestrations into the pore (‘full’ lipid block, *Fig. 1C*), which occurred in one equilibrium replica of Ca^2+^-free (*SI Fig. 1A*; in the simulations at +300 mV spawned from this replica lipids remained in the pore), as well as in the two events of spontaneous dehydration of the pore in Ca^2+^-bound at +300 mV, described in more detail below. In terms of the channel’s conductance, in simulations with applied voltage Ca^2+^-bound showed steady outward K^+^ permeation as long as the pore remained hydrated and not blocked by lipids (24±11 pS), and Ca^2+^-free didn’t permeate as long as either partial or full block occurred. In terms of pore hydration, consistent presence of at least 20 water molecules in the pore (using our definition of the pore region; *see Methods*) appeared to correlate with a conductive channel. It should be noted that the observed conductance value is an order of magnitude lower than the experimental one (up to 300 pS) ^4^ - a known feature of force fields without explicit description of polarization, such as the commonly used in biomolecular simulations CHARMM36m, also employed in this study ^16,44^. Notably, in those simulations of the Ca^2+^-free state at +300 mV where the pore remained hydrated and not blocked by lipids, the channel showed steady outward K^+^ permeation of 8±3 pS (*Fig. 1D, SI Fig. 1C, D, SI Table 1*). In contrast to this observation, experiments show similar single-channel conductances of BK in both presence and absence of Ca^2+ 45^. This discrepancy could be explained by the channel primarily exploring the conformations close to the initial structure (i.e. mostly functionally closed when initiated from the Ca^2+^-free, and mostly functionally open when initiated from the Ca^2+^-bound structure) on the MD timescales.

**Fig. 2.**
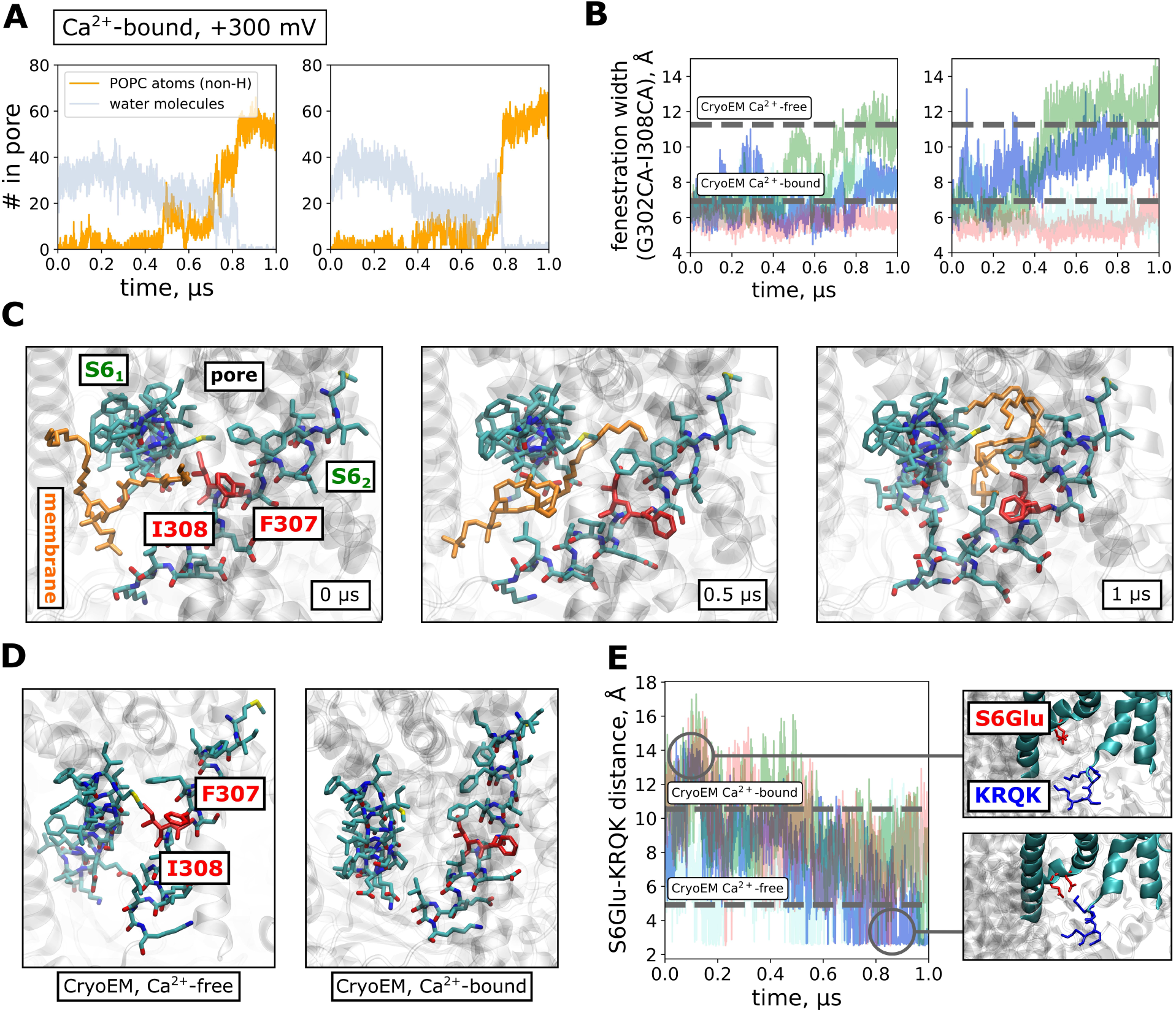
Simulation replicas of Ca^2+^-bound at +300 mV where spontaneous lipid block occurred. (*A*) Time traces of lipid entry and pore dehydration. (*B*) Time traces of fenestration width, defined as the distance between CA atoms of G302 residues of opposing subunits. (*C*) Simulation snapshots showing lipid entry into the pore and the accompanying opening of a fenestration between the pore-lining S6 helices, with motions of residues I308 and F307 (colored red). (*D*) CryoEM structures of Ca^2+^-free and Ca^2+^-bound showing conformational differences similar to those observed during the spontaneous lipid block of Ca^2+^-bound. (*E*) Formation of salt bridges between E310/E310 and ^317^KRQK^320^ during the spontaneous lipid block, shown for one of the simulation replicas of Ca^2+^-bound where the spontaneous lipid block occurred, indicative of an open to closed state transition; different colors represent different E310/E313-KRQK pairs (four in total).

**Fig. 3.**
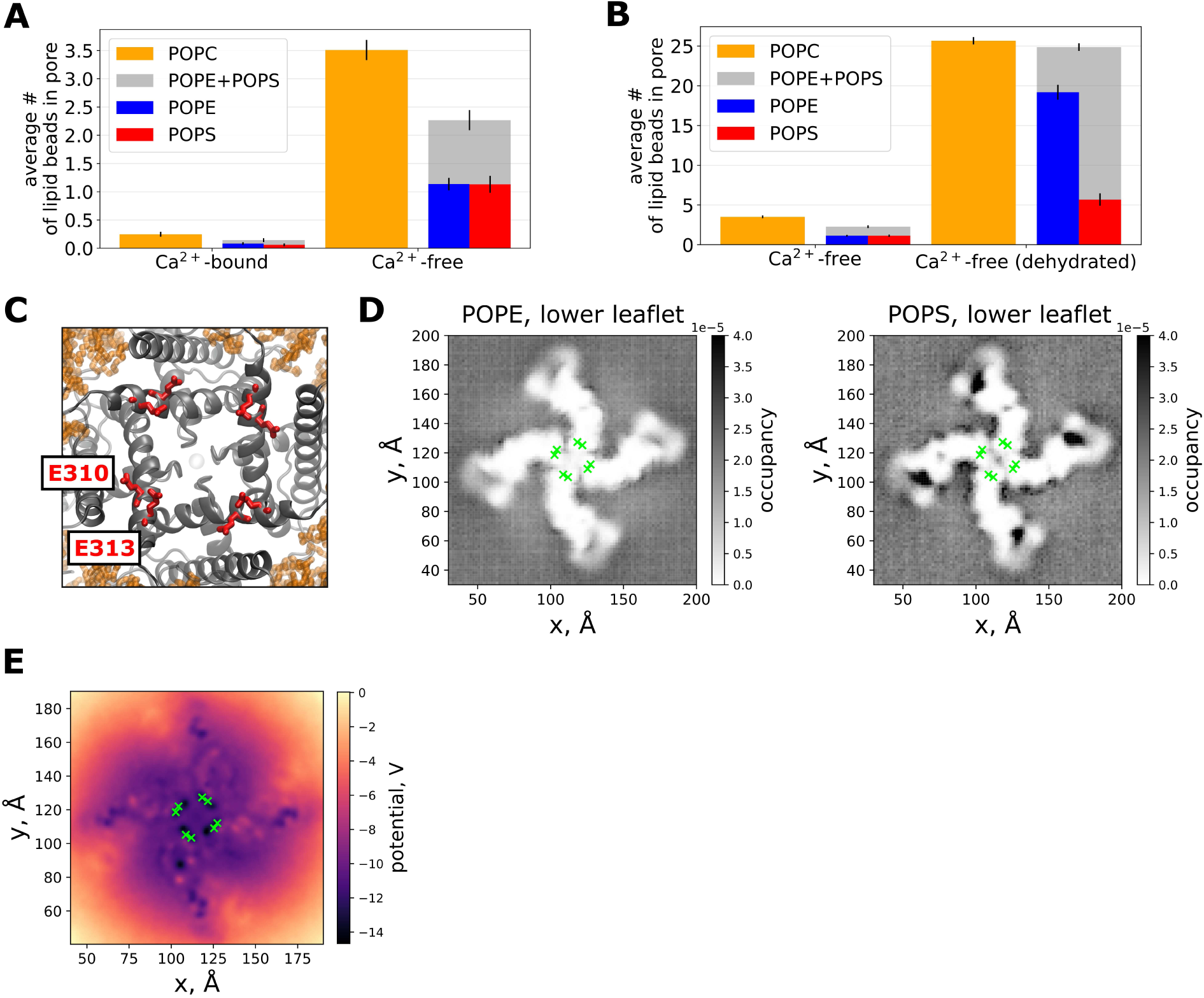
(*A*) Lipid entry in Ca^2+^-bound and Ca^2+^-free states, in POPC and POPE:POPS membranes in Martini3 simulations with a hydrated pore in both systems, and (*B*) with restraints preventing water beads from entering the pore in the Ca^2+^-free state, shown on the right, and the Ca^2+^-free state with a hydrated pore shown again for comparison (left). (*C*) E310/E313 residues on the pore-lining S6 helices, (*D*) occupancies of POPE (*left*) and POPS (right) headgroups around the channel with E310/E313 depicted in green (Ca^2+^-free state is shown), and (*E*) average electrostatic potential in Martini3 simulations at the level of E310/E313.

Such diverse behavior of the Ca^2+^-free state - a hypothetical closed state of the channel - in terms of lipid block and conductance, raises the question of the relationship between pore conformation and the conductive state of the channel. Generally, we observed a variety of pore conformations in the simulations of the Ca^2+^-free state. These conformations often differed from the initial CryoEM structure, which specifically manifested in F304 side chains rotating into the pore, as well as overall pore narrowing (*Fig. 1E, F, SI Fig. 2*) - similar to observations in an early computational study ^34^, and resembling the relative pore narrowing in a recent CryoEM structure of Ca2+-free BK with a deactivating mutation in the voltage sensor, proposed to represent a deeper closed state of the channel ^46^. To note, in one simulation replica without applied voltage, the pore became effectively sealed at the residue I308, indicating that full pore closure can in principle be achieved (*SI Fig. 2A, ‘rep1’*) - however, this was concurrent with lipids occupying the pore (*SI Fig. 1A, ‘rep1’)*, thus it is unclear whether full pore closure (and thus a possible steric block of ion permeation posed by the channel itself) is possible without lipid block. Overall, the conformational changes in the Ca^2+^-free state showed some degree of correlation with the conductive state of the channel; for instance, the aforementioned simulation replicas of Ca^2+^-free under voltage which did not undergo pore dehydration/lipid block and were conductive, showed a smaller degree of pore narrowing and F304 rotation relative to the CryoEM structure compared to the replicas which did undergo pore dehydration/lipid block and were not conductive (*SI Fig. 2B*). To note however, the equilibrium replica which underwent full lipid block, together with the simulations under voltage spawned from this replica (where the channel remained to be blocked by lipids and was not conductive), did not show deviation in the pore width compared to the Ca^2+^-free CryoEM value - and showed a smaller degree of F304 rotation (see *SI Fig. 2A, ‘rep4’* for the behavior of the equilibrium replica, and *SI Fig. 2C* for the consecutive replicas under applied voltage). This might indicate compatibility of the Ca^2+^-free pore’s dimensions in the CryoEM structure with full lipid block, and suggest dependence of conformational behavior on whether lipid block is partial or full. Thus, to summarize, based on our equilibrium and applied voltage simulations of the full-length BK, we confirm that the Ca^2+^-bound structure corresponds to an open, conductive state of BK. At the same time, the interplay between further conformational changes in the channel’s pore relative to the Ca^2+^-free CryoEM structure and the obligatory pore dehydration/lipid block was needed to stop ion permeation in the Ca^2+^-free state.

### Spontaneous closure of the open state

As we mentioned earlier, the Ca^2+^-bound state remained hydrated and conductive throughout most of our simulations; however, in two out of six replicas at +300 mV the pore of the open channel eventually underwent spontaneous dehydration, accompanied by a full lipid block of the pore in both cases (*Fig. 2A*; all Ca^2+^-bound +300 mV replicas including those where lipid block didn’t happen are shown on *SI Fig. 3* for comparison) - in addition to one equilibrium replica with decreased pore hydration and increased lipid presence in the pore (*SI Fig. 1B, ‘rep2’*). To note, spontaneous unbinding of the activatory Ca^2+^ from its CryoEM binding sites was a consistent phenomenon throughout the simulations. Unbinding occurred predominantly from the RCK1 site, with one ion unbinding during equilibrium simulations with more following in applied voltage simulations (*SI Fig. 5*), which could contribute to the reduced structural stability of the Ca^2+^-bound state. Generally, the entrance of an entire lipid molecule was made possible by widening of one of the fenestrations between S6 helices of neighboring subunits (*Fig. 2B;* all replicas are shown on *SI Fig. 4B* for comparison), accompanied by motions of bulky hydrophobic residues obscuring the fenestrations in the Ca^2+^-bound structure, such as F307 and I308 (*Fig. 2C*). Interestingly, opening of the fenestrations and motions of these residues strongly resemble the respective differences between Ca^2+^-bound and Ca^2+^-free CryoEM structures, supporting the notion that we observed transitions from the open to the closed state of the pore (*Fig. 2D*). Additionally, in a previous study that combined MD simulations and mutagenesis experiments, the positively charged residue clusters at the end of S6 helices (corresponding to ^317^KRQK^320^ in the BK from *Aplysia californica* studied here) and S6 glutamates of the neighboring subunits (‘S6Glu’) were shown to play a role in the stabilization of the open state of the channel, and proposed to form salt bridges in the closed, but not the open state of the channel ^24^. In our simulations, we observed clear formation of such salt bridges before fenestration widening in one of the two replicas with a spontaneous lipid block (*Fig. 2E*). In the other replica with a full lipid block, the salt bridges were forming only transiently - similar to several other replicas with no lipid block; additionally, one of the replicas with no lipid block showed formation of a relatively long-lived salt bridge (*SI Fig. 4A*). Taken together, this could indicate the existence of mechanisms that induce fenestration widening (and thus increased propensity for a lipid block) with some degree of independence of salt bridge formation - or, alternatively, taking into account Ca^2+^ unbinding in all replicas, the possibility of pore closure and lipid block happening at longer simulation times in these replicas as well.

**Fig. 4.**
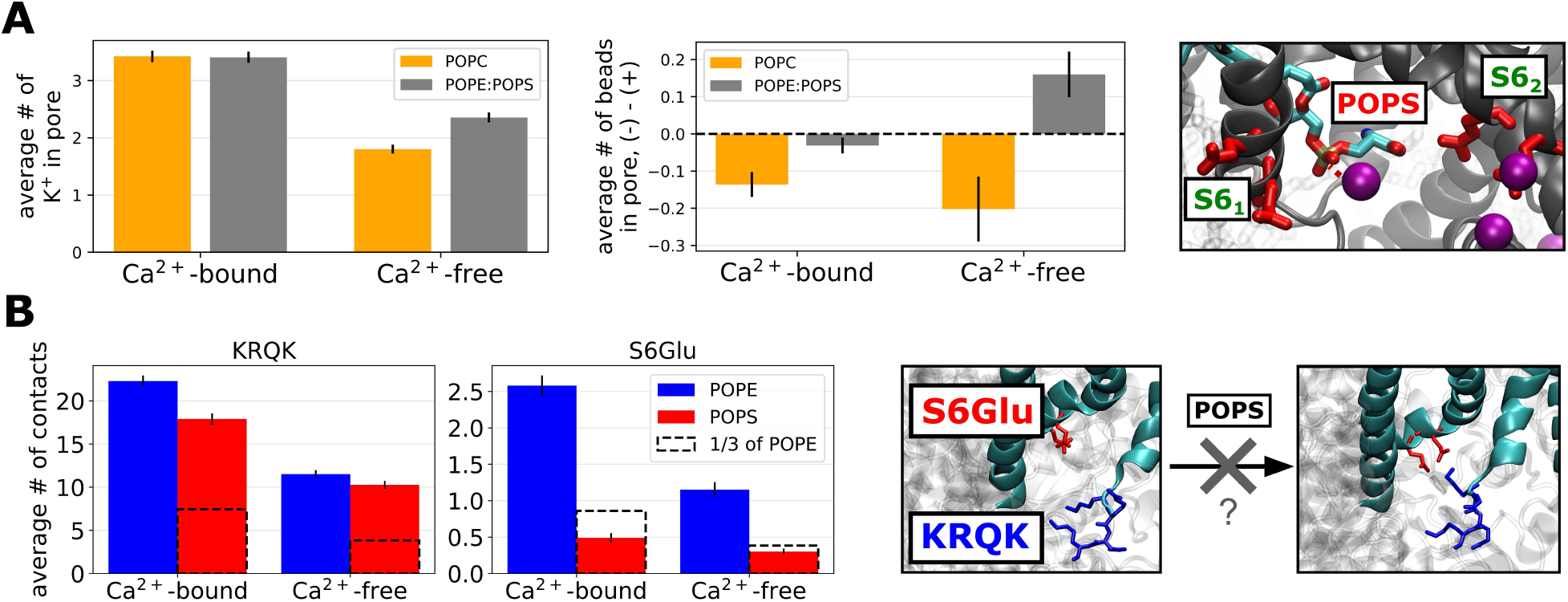
(*A*) Difference in pore occupancy by negatively charged beads and positively charged beads in Martini3, showing increase in the average number of K^+^ beads in pore in the Ca^2+^-free state with a hydrated pore (*left*), accompanied by enrichment of the pore by negatively charged beads for that system in a POPE:POPS membrane (*middle*), explained by electrostatic attraction of K^+^ into the pore by negatively charged headgroups of POPS transiently occupying the fenestrations between S6 helices and the pore; E310/E313 residues are shown for reference (*right*). (*B*) Average number of contacts of each lipid type with either the positively charged ^317^KRQK^320^ cluster, or the negatively charged E310/E313 (‘S6Glu’, *left*). ‘⅓ of POPE’ shows the hypothetical number of POPS contacts if the POPS contact frequency was exactly proportional to its fraction in the membrane relative to POPE (⅓) - indicating that POPS disproportionately frequently binds to KRQK and is repelled from the E310/E313, thus possibly perturbing the formation of a salt bridge between ^317^KRQK^320^ and E310/E313 characteristic of the closed state.

**Fig. 5.**
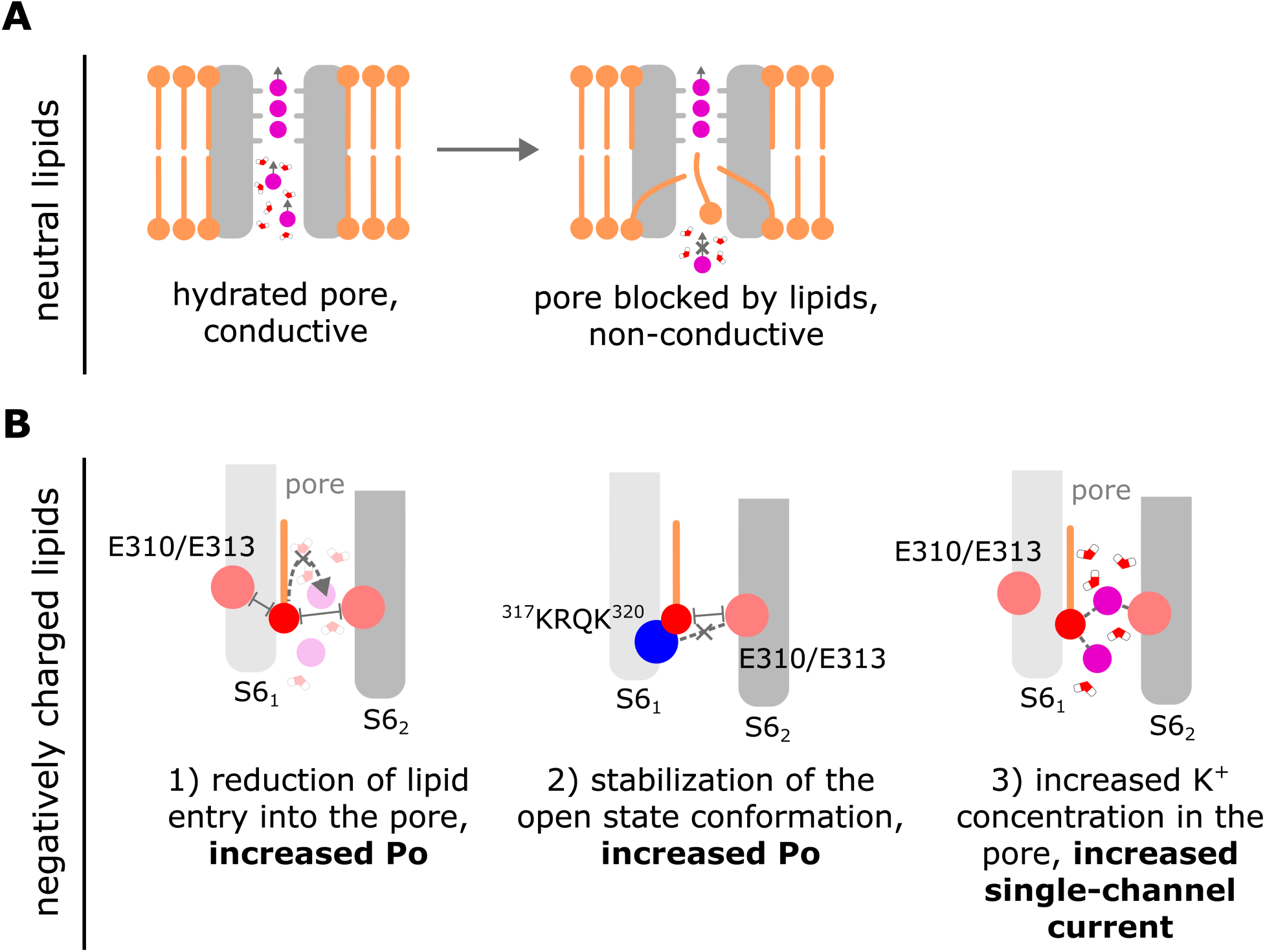
A scheme illustrating the main hypotheses of this work. (*A*) While the pore of the channel in the open, conductive state of the channel remains hydrated, the closed state is characterized by either lipid tails, or entire lipid molecules entering the pore through membrane-facing fenestrations and blocking ion permeation. (*B*) Presence of negatively charged lipids in the membrane leads to 1) reduced lipid entry into the pore due to the electrostatic repulsion from the fenestration-lining glutamate residues in S6 helices, 2) increased K^+^ concentration in the pore and thus increased probability of a permeation event due to the electrostatic attraction between K^+^ and negatively charged headgroups spontaneously occupying the fenestrations and the pore, and 3) disruption of the ^317^KRQK^320^–E310/E313 salt bridge formation, proposed to be a crucial step in channel closure ^24^, via interaction of the negatively charged lipids with ^317^KRQK^320^ and repulsion from E310/E313. These effects explain the experimentally observed increase in the open probability and the single channel-current in negatively charged lipid-containing membranes ^42,43^.

### Effect of membrane composition on lipid block

Membrane lipid composition plays a significant role in regulating the behavior of BK channels ^47,48^. Here, we focus on the negatively charged POPS lipid, the presence of which - and negatively charged lipids in general - was shown to increase both the open probability (Po) of the channel and its single-channel current ^43^. To tackle this issue computationally, we first performed coarse-grained simulations of Ca^2+^-bound and Ca^2+^-free states of BK using the Martini3 force field ^49^ in a POPE:POPS (3:1 mol.) membrane - the ratio used in the experimental study ^43^ - with control simulations in a POPC membrane (*SI Table 2*). The switch from all-atom to coarse-grained simulations was done to increase the accessible simulation times and extensively sample the relevant configurations of lipids around the protein. After 40 µs-long simulations for each combination of the channel state and membrane composition, we observed a significant decrease in the pore occupancy by lipids in POPE:POPS compared to POPC (*Fig. 3A*). Interestingly, the pore stayed hydrated in the Ca^2+^-free state (*SI Fig. 6A*). To note, in the Martini3 variant with elastic networks that we used ^49,50^, the conformational changes of the Ca^2+^-free we observed in the atomistic simulations were restricted, which might have increased the barrier for pore dehydration and/or lipid entry, reflecting the previous observation of a certain degree of correlation between the propensity for lipid block and further conformational changes relative to the Ca^2+^-free CryoEM structure, including the simulation replicas where Ca^2+^-free remained hydrated and conductive. In line with the hydrophilic environment of the pore, the number of the Martini lipid beads in the pore in Ca^2+^-free was comparatively low for both POPE:POPS and POPC membranes - on average ∼2-3 out of 12 beads in a single residue for every lipid type, which would correspond to ∼8-12 out of ∼50 non-hydrogen atoms in a single lipid in atomistic representation, in contrast to ∼20 or more for partial lipid block in atomistic simulations (*SI Fig. 1D*). Reflective of the hydrophilic environment, it was lipid headgroups which occupied the pore most frequently as opposed to tails (*SI Fig. 7*). When water was artificially excluded from the pore of the Ca^2+^-free state using restraints, POPE:POPS and POPC membranes showed practically no difference in lipid entry, with a drastically increased average number of lipid beads in the pore for both membrane compositions (*Fig. 3B, SI Fig. 6B*). Given the absence of the effect of POPS on the fully closed state characterized by a hydrophobic pore and substantial lipid entry, this suggests that the presence of POPS in the membrane increases the barrier for lipid entry in open and intermediate closed states, as long as the pore is still hydrated.

Considering the possible mechanistic reason for this increased barrier for lipid entry, interestingly, not only did the average number of lipid beads in the pore decrease in presence of POPS, but so did the average number of individual lipid residues simultaneously occupying the pore (*SI Fig. 7C*). Notably, the S6 helices contain two negatively charged glutamates per chain - E310 and E313, the same glutamates that form salt bridges in the closed state with the KRQK cluster at the lower ends of S6 helices mentioned above (*Fig. 3C*). Here however, the presence of eight negatively charged residues close to the fenestrations between S6 helices could in principle provide a barrier for entry of similarly charged POPS molecules. This would be especially relevant for lipid entry into the pore, as even though POPS comprises only a quarter of the membrane, the area around the protein is actually enriched with POPS, including the region near the pore (*Fig. 3D, SI Fig. 8A*). Indeed, the average electrostatic potential calculated at the level of E313, which coincides to a large extent with the position of lipid headgroups near fenestrations (*SI Fig. 8B*), is more negative both in the fenestrations and the pore relative to the regions of the membrane close to the fenestrations in Martini simulations (*Fig. 3E*). Thus, we suggest the electrostatic repulsion between E310/E313 and the negatively charged POPS headgroups as the driving force behind reduced lipid entry, explaining the increased open probability of the channel in presence of POPS observed in experiments ^42,43^.

Previously, we mentioned that in simulations of Ca^2+^-bound and Ca^2+^-free states with a hydrated pore it was primarily the headgroups that contributed to the transient occupancy of the pore by lipids. Aligning with the larger POPS occupancies in the membrane regions surrounding the protein, the pore in POPE:POPS was enriched by negatively charged beads (measured as the occupancy difference between the negatively and positively charged beads), compared to POPC in our Martini3 simulations, with the biggest increase in the simulations of the Ca^2+^-free state with a hydrated pore (*Fig. 4A, middle*). In the Ca^2+^-bound state this effect was less pronounced, likely due to the tightly closed fenestrations (*Fig. 4A, middle*). At the same time, in the Ca^2+^-free state with a hydrated pore the average number of K^+^ residing in the pore increased, whereas it remained practically unchanged in the Ca^2+^-bound state (*Fig. 4A, left*). It is possible that a similar increase in local K^+^ concentration in the pore could generally happen in open, Ca^2+^-bound-like conformations due to the channel’s intrinsic flexibility (and e.g. some degree of fenestration opening that would allow the negatively charged headgroups to be closer to the pore), which was however restricted by the elastic networks in the Martini force field. In our control atomistic simulations of Ca^2+^-bound in a POPE:POPS membrane, we did observe an increase in the average number of K^+^ ions in the pore in one of the two POPE:POPS simulation sets, compared to the other with a different initial membrane configuration (as well as relative to POPC simulations) - while POPS headgroups were present in the pore in this simulation set and absent in the other. It should be noted that the number of K^+^ ions in the pore correlated with the pore occupancy by lipid tails as well, which was different in both POPE:POPS simulation sets, as well as in POPC, possibly reflecting the limited sampling in the atomistic simulations (*SI Fig. 9*). Taking these force field- and sampling-related limitations into account, and drawing from the mechanism of activation proposed for small-molecule negatively charged activators of K^+^ channels (NCAs) ^51^, we cautiously propose the explanation for the increased single channel current by negatively charged lipids: the headgroups of negatively charged lipids occupy the fenestrations (or transiently the pore), thereby electrostatically attracting and increasing the local concentration of K^+^ in the pore, leading to an increased probability of a permeation event, via multi-ion knock-on mechanism (*Fig. 4A, right*).

The effects we observed so far relate primarily to the propensity of different lipids to enter the pore, which can be both inhibitory, as is the case for lipid block, and activatory, in case of the transient pore occupancy by POPS headgroups. However, another possible dimension of the activatory effect of negatively charged lipids is stabilization of the open state conformation of the channel. In Martini simulations we observed disproportionately frequent interaction of POPS (i.e. with a frequency bigger than the fraction of POPS in the membrane, or 1/4) with the KRQK cluster, and, conversely, rare interactions with the S6 glutamates, reflective of the electrostatic attraction between POPS molecules and KRQK, and repulsion from E310/E313 (*Fig. 4B*). This, in principle, could interfere with the formation of the S6 glutamates-KRQK salt bridge characteristic of the closed state, an effect proposed recently for the by phosphatidylinositol 4,5-bisphosphate (PIP2, charge -3) activation of the SK2 channel, in which similar salt bridges can form ^52^. To evaluate these conformational effects, we performed control atomistic simulations of BK in POPE:POPS (*SI Table 2*). We observed a higher frequency of states with larger KRQK-glutamate distances compared to the simulations in POPC, indicating stabilization of the pore domain’s structure (*SI Fig. 10A*). Similar to the Martini simulations, S6Glu was depleted of POPS relative to POPE in both simulation sets, and KRQK enriched with POPS in one simulation set; in the other set, KRQK did not show disproportionately frequent interactions with POPS relative to its molar fraction in the membrane, nevertheless the existing POPS-KRQK contacts and the repulsion of POPS from S6Glu appeared to be sufficient to stabilize the channel’s conformation in this case (*SI Fig. 10B*). Additionally, we performed atomistic simulations of an isolated pore domain in POPE:POPS and POPC as an approach to increase sampling due to a largely reduced size of the system (*SI Table 2*). However, despite longer simulations and a larger number of initial membrane configurations tested, the isolated pore domain showed very low conformational stability in both membrane compositions (*SI Fig. 11*). In summary, based on our simulations we propose a multi-modal mechanism of activation by negatively charged lipids, involving reduced lipid entry, increased K^+^ occupancy of the pore and stabilization of the open state conformation. To note, in our Martini3 simulations POPS also showed distinct binding sites in the outer voltage sensor region (*SI Fig. 12*), which might be related to the inhibitory effect of another negatively charged lipid (PIP2) observed experimentally at high voltages ^53^ (see *Discussion*), although a detailed investigation of this effect is beyond the scope of our study.

## Discussion

BK channels present an interesting case where CryoEM structures, despite being obtained in activating and deactivating conditions, do not provide a clear definition of open (conductive) and closed (non-conductive) states, particularly due to the absence of a physical constriction in the permeation pathway in deactivating conditions, such as a helix-bundle crossing ^54,55^, or a collapsed SF ^56,57^ occurring in some other K^+^ channels. Later, computational and experimental studies suggested a model where the channel’s pore in the closed state undergoes dehydration, thus establishing a hydrophobic barrier to permeation of K^+ 34–36,39^. However, to our knowledge, the ion permeability of the BK CryoEM states has not been tested directly in the full length BK, such as with MD simulations with applied voltage. At the same time, the pore-lining S6 helices of neighboring subunits form membrane-exposed fenestrations in Ca^2+^-free structures (deactivating conditions), which in some CryoEM structures contain lipid tails ^11^, in addition to spontaneous occupancy of the pore by lipid tails in MD studies ^34^, shown to increase the hydrophobicity of the pore in the closed state in a recent computational study ^40^. In this work, we focused on the role of lipids in the BK channel’s gating. Our MD simulations of the full-length channel in equilibrium and under applied voltage allowed us to test the conductivity of the full length channel in Ca^2+^-bound and Ca^2+^-free states directly, and show that the pore block by lipids is a defining characteristic of the closed, non-conductive state of the channel. Pore block occurred in two modes: in one mode, only tails of lipids can enter the pore through the S6 fenestrations, likely not only increasing the hydrophobicity of a dewetted closed-state pore, but also posing a steric barrier to ion permeation (‘partial block’, *Fig. 1C, left*). Second, in certain cases (including the events of spontaneous pore closure in the Ca^2+^-bound state) entire lipid molecules were able to enter into the pore through the fenestrations (‘full block’, *Fig. 1C, right*). Both modes of lipid block were able to stop ion permeation on MD timescales (microseconds). Generally, both mechanisms align with experimental data, including some of the CryoEM structures of BK containing lipid tails in fenestrations mentioned earlier (supporting partial block) ^11^, mutants of F315 (human BK, equivalent to F304 in *Aplysia* BK) located in fenestrations having activatory/deactivatory effect depending on their size (bigger residues activate, while smaller deactivate the channel) ^58,59^, possibly indicating modulation of the steric barrier for lipid entry, as well as the dependence of Po and single-channel currents on the membrane composition (such as activation of the channel by lipids with bigger headgroups, which can be explained by a higher energetic barrier for full lipid entry through the fenestrations between the S6 helices) ^47,48,60^. To note, BK in the fully closed state was shown to be insensitive to the large pore blocker bbTBA ^19^ - while we don’t preclude that there might be states where the pore is completely inaccessible to bbTBA due to sterical barrier provided by e.g. full lipid block (the pore in WT Ca^2+^-free CryoEM states is wide enough to fit bbTBA), a recent CryoEM structure of BK with a deactivating mutation in the voltage sensing domain showed the pore narrow enough to prevent bbTBA entry, which could also explain lack of closed-state block by bbTBA ^46^. At the same time, in a CryoEM structure of a K^+^ channel from the K2P family (TREK-1), which has similar membrane-exposed fenestrations in the closed state, it was possible to resolve a lipid molecule occupying the pore ^61^ A recent CryoEM study also showed lipid densities both in fenestrations and inside the pore of another K2P family member TWIK-1 ^62^, which not only confirms the results of an earlier MD study showing lipid entry into the pore in TWIK-1 ^63^ - but also indicates that full lipid block can generally serve as a pore closure mechanism - albeit further structural studies would be needed to confirm that for BK channels.

The observation of two modes of lipid block - partial and full - also raises a question, whether they could correspond to different functional states of BK. In electrophysiological recordings, BK can occupy several open (conductive) and closed (non-conductive) states that have different life times ^64^. Naively it can be assumed that the barrier to relieve the partial block (move only the lipid tails out of the pore) is lower than that for the full block, thus in the framework of lipid block as the closing mechanism, the partial block would correspond to the short-lived closed states, and full block to the long periods of closure.

One hypothesis regarding the Ca^2+^-free state lacking a physical constriction of the pore is that it does not represent the final closed state, but rather an intermediate one, and further closure of the pore is possible ^8^, as is the case for a prokaryotic homolog of the BK channel, MthK, which shows a helix bundle-like structure in its closed state ^65^. Indeed, later cryoEM structures and MD simulations of BK with mutations in the voltage sensor allowed to capture a state with a narrower pore (albeit wider than a hydrated K^+^ ion) ^12,34^. Further pore narrowing also occurred in practically all of our simulations of the Ca^2+^-free state where the partial lipid block happened - both relative to the CryoEM structure and the atomistic Ca^2+^-free simulations where the pore remained hydrated. Additionally, in Martini3 simulations where such changes were limited by elastic networks, the pore remained hydrated and with a relatively low degree of lipid entry in the Ca^2+^-free state, unless restraints preventing pore hydration were used, presenting further evidence of the importance of additional conformational changes in achieving the fully closed state. However, it should be noted that the mode of pore closure here differed from the helix bundle mechanism - the pore closed only partially in all replicas except one with full pore closure accompanied by lipid block - and at least in all cases we observed it appeared to be only accessory to lipid block.

Notably, pore wetting has been studied for model non-biological nanopores, where the hydration state of the pore appears to depend on its radius and geometry; moreover, for a pore of a given that is dehydrated at 0 V, wetting can be induced by applied voltage ^66^. Such voltage-dependent behavior was observed in K^+^ channels as well, such as the negative hydrophobic pore collapse in the Kv1.2 channel induced by negative voltages ^67^. While a scan of various voltages and their effect on the dynamics of lipids/water in the BK channel’s pore is beyond the scope of this work, this presents an interesting direction for future research, especially considering the conformational flexibility of BK’s pore and lipids contributing to pore hydrophobicity.

Finally, we investigated the mechanism of BK activation by negatively charged lipids. It has been shown that such lipids can increase both the Po of the channel, as well as single-channel current ^42,43^. In light of our observation of lipid block characterizing the closed state of the channel, it was interesting to test whether activation by negatively charged lipids is related to different lipids having different propensities for lipid block. Based on our atomistic and coarse-grained simulations of a membrane containing the negatively charged POPS lipid, we suggest a multi-modal mechanism of its action, with 1) reduced lipid entry in open and intermediate closed states when the pore is still hydrated, and thus increased Po, 2) increased local K^+^ concentration in the pore, which explains the increase in the single-channel current, and 3) stabilization of the open-state conformation of the channel via disruption of the ^317^KRQK^320^– E310/E313 salt bridge formation, also explaining the increased Po. Considering the latter mechanism, a mutagenesis study featuring another negatively charged lipid - PIP2 - showed that in the human BK channel, it interacts specifically with the RKK motif, equivalent to KRQK in the *Aplysia* BK ^42^. Additionally, for another K^+^ channel with similar positively/negatively charged residues in the lower S6 helices - SK2 - MD simulations and mutagenesis also suggested disruption of the salt bridge and activation by PIP2 ^52^. Even though experimentally this effect was studied specifically for PIP2, we hypothesize that the channel’s open-state structure might be stabilized via this mechanism by negatively charged lipids in general.

In terms of the physical basis of activation by negatively charged lipids, electrostatic interactions played a key role: eight E310/E313 residues on S6 helices contribute to regions of negative electrostatic potential near and in the pore that increases the barrier for POPS molecules to enter it (*Fig. 3C, E*). At the same time, the general enrichment of the areas near the channel by POPS enhances this effect on lipid entry, despite POPS constituting only 25% of the membrane in our case (*Fig. 3D, SI Fig. 8A*). In experiments, the open probability of the channel was shown to be higher with a bigger negative charge on the lipid’s headgroup ^42^, which is in agreement with the idea of electrostatic repulsion of the negative charges on the lipids from the pore by S6 glutamates. Conversely, positively charged lipids were shown to decrease the open probability^68^ - in the framework of the model we propose, their effect could be explained by negatively charged E310/E313 attracting the positively charged lipids into the pore, although this would require further validation (such as by studying the effect of E310/E313 mutants on the channel’s activity in positively and negatively charged lipid-containing membranes; it should be noted that such mutants might affect not only lipid entry, but also the conformational stability of open and closed states, given the role that E310/E313 also play in the latter ^24^). Similarly, POPS headgroups residing close to the pore attracted K^+^ into it (*Fig. 4A*) - we propose that the electrostatic attraction between the negatively charged headgroups and K^+^ lies behind the increase in the single channel current, akin to the mechanism proposed for small-molecule negatively charged activators of K^+^ channels ^51^. In addition, an earlier experimental study showed POPS-containing bilayers altering the channel’s conductance, but not Ba^2+^ and tetraethylammonium block ^69^: this could be explained by the additional negative electrostatic potential provided by POPS being cancelled out by a locally increased K^+^ concentration in the pore.

Interestingly, some previous studies suggested a dual effect of the negatively charged PIP2 - not only activatory, but also inhibitory at high enough voltages ^53^. Such inhibition was hypothesized to originate from PIP2 binding to an intersubunit cleft between two voltage sensors, thereby augmenting the negative charge on the cytosolic side of the membrane and effectively pulling the voltage sensors into the ‘down’ (inactivated) configuration, as evidenced by suppression of inhibition in presence of voltage-sensor-binding β subunits, or via shielding of charges by intracellular divalent cations ^53^. In our Martini3 simulations, POPS forms a distinct binding site in the outer voltage sensor region (*Fig. 3D, SI Fig. 8A, SI Fig. 12*). This generally supports the notion of involvement of negatively charged lipids in voltage activation, however the associated conformational changes in the voltage sensor and their coupling to the pore domain are beyond the scope of this work.

To conclude, we present a model of BK gating where lipids play an integral role in determining the conductive state of the channel, and suggest a multi-modal mechanism of how the negative charge of lipid headgroups can activate the channel (*Fig. 5*). BK channels exist in complex multicomponent lipid environments, such as plasma, mitochondrial and nuclear membranes, also in various cell types ^7^. At the same time, the BK channel activity was shown to be regulated, in addition to the lipid headgroup charge, by headgroup size, tail length, as well as specific protein-lipid interactions ^41–43,47,48,60,68^. This likely leads to intricate patterns of regulation in multicomponent membranes, not only presenting further opportunities to study and validate the role of lipids in the function of BK channels, but also help inspire the design of similarly acting new small molecule modulators of BK channels.

## Methods

### System preparation

CryoEM structures of BK from *Aplysia californica* in Ca^2+^-bound (PDB ID 5TJ6) and Ca^2+^-free (PDB ID 5TJI) states ^8,9^ were used throughout this study. To build the all-atom systems, the CryoEM structures of the full length channel were embedded in a 1-palmitoyl-2-oleoyl-glycero-3-phosphocholine (POPC) lipid bilayer using CHARMM-GUI ^70,71^. The non-terminal loops missing from the structures were added, and N- and C-termini were acetylated and methylamidated, respectively. The system with the Ca^2+^-bound structure contained 1523 POPC molecules, 3641 K^+^, 3649 Cl^-^, 247063 water molecules, as well as 4 Mg^2+^ and 8 Ca^2+^ ions in their CryoEM binding sites. The system with the Ca^2+^-free contained 1503 POPC, 3629 K^+^, 3609 Cl^-^ and 245986 water molecules. This includes 4 K^+^ added to the SF (binding sites S1-S4), as the CryoEM structure of the Ca^2+^-free state, opposite to Ca^2+^-bound, doesn’t have any K^+^ resolved inside the SF. Additionally, 2 water molecules per subunit were added to the cavities behind the SF for each CryoEM structure, as water molecules in these positions are usually present in high-resolution structures of other channels, and were shown to have a stabilizing effect on the SF structure ^72–75^. The number of K^+^ ions was chosen to yield a concentration of 0.8 M, and the number of Cl^-^ to yield systems with a total charge of zero.

The coarse-grained structures of the protein for Martini3 ^49^ simulations were obtained using martinize2 ^76^, and the protein-membrane systems were built using insane ^77^. For the simulations with POPC membranes, the system with the channel in the Ca^2+^-bound state contained 1602 POPC molecules, 3400 K^+^ beads, 3384 Cl^-^ beads and 53638 water beads (regular); the system with the channel in the Ca^2+^-free state contained 1546 POPC molecules, 3407 K^+^, 3387 Cl^-^ and 53722 water beads. The systems with 1-palmitoyl-2-oleoyl-sn-glycero-3-phosphoethanolamine (POPE) and 1-palmitoyl-2-oleoyl-sn-glycero-3-phospho-L-serine (POPS) and a 3:1 ratio contained, in case of the Ca^2+^-bound state, 400 POPS, 1201 POPE molecules, and 3600 K^+^, 3184 Cl^-^ and 53642 water beads; in case of the Ca^2+^-free state, 385 POPS, 1159 POPE molecules, 3599 K^+^, 3194 Cl^-^ and 53716 water beads.

Systems containing the full-length channel for the control atomistic simulations in the POPE:POPS membrane were built similarly to the simulations of the full-length channel in POPC, as was the case for the isolated pore domain in POPE:POPS and POPC. To mitigate the limited membrane configuration sampling due to slower lipid dynamics compared to the Martini3 simulations ^49^, two systems with different initial configurations of lipids in the membrane were built for each state of the full-length channel in POPE:POPS. For the Ca^2+^-bound state of the protein, one system contained 1200 POPE, 400 POPS, 3367 K^+^, 2975 Cl^-^, 226858 water molecules, as well as 4 Mg^2+^ and 8 Ca^2+^ in their binding sites; the other contained 1200 POPE, 400 POPS, 3368 K^+^, 2976 Cl^-^, 226877 water molecules, 4 Mg^2+^ and 8 Ca^2+^. For the Ca^2+^-free state, one system contained 1125 POPE, 375 POPS, 3191 K^+^, 2796 Cl^-^ and 215037 water molecules; the other contained 1125 POPE, 375 POPS, 3191 K^+^, 2796 Cl^-^ and 215033 water molecules.

For the isolated pore domain, one system in the POPC membrane, and four systems in the POPE:POPS membrane with different initial configurations of lipids in the membrane were built for each state of the channel. The POPC system with the Ca^2+^-bound state contained 127 POPC, 106 K^+^, 94 Cl^-^ and 7167 water molecules; with Ca^2+^-free 126 POPC, 114 K^+^, 102 Cl^-^ and 7676 water molecules. In POPE:POPS, for the Ca^2+^-bound state each system contained 99 POPE, 33 POPS and, correspondingly, 142 K^+^, 97 Cl^-^, 9644 water molecules for systems 1 and 2; 142 K^+^, 97 Cl^-^, 9619 water molecules for system 3; and 141 K^+^, 96 Cl^-^, 9613 water molecules for system 4. For the Ca^2+^-free state, they contained 99 POPE, 33 POPS, 147 K^+^, 102 Cl^-^ and 1) 9969, 2) 9973, 3) 9979, 4) 9959 water molecules.

### MD simulations

The systems containing the full-length channel were first equilibrated using the default 6-step protocol with the gradual release of restraints on the protein and lipids provided by CHARMM-GUI ^70,71^ to reach the temperature of 323 K and the pressure of 1 bar. CHARMM36m/TIP3P parameters were used for all atomistic simulations in this work. All simulations throughout this work were carried out using GROMACS, versions 2020 to 2024 ^78^. Simulations without applied voltage were then run in the NPT ensemble using the semi-isotropic Parrinello-Rahman barostat^79^ and velocity rescale (v-rescale) thermostat ^80^. Periodic boundary conditions (xyz) were used. Hydrogen-containing bonds were constrained using the LINCS algorithm ^81^; simulation timestep was 2 fs. Lennard-Jones interactions were treated using a cutoff scheme, with forces switched to 0 between 1.0 nm and 1.2 nm. Coulomb interactions were treated using PME ^82^ with a 1.2 nm cutoff. Five simulation replicas were run for each system without voltage. Simulations with applied voltage of the full-length channel in POPC were started from frames of equilibrium simulation replicas at 0.2-0.5 µs, isolating the functional state of interest (full, partial or no lipid block), and run for up to 1 µs/replica, 3-6 replicas per system. To reach the desired transmembrane voltage *ΔV* of +300 mV, a constant electric field was applied, calculated as *E = ΔV/dz*, where *dz* is the size of the box along the *z-*axis ^83^. All simulations done for the *Results* subsections *All-atom simulations of Ca^2+^-bound and Ca^2+^-free states* and *Spontaneous closure of the open state* are listed in *SI Table 1*; simulations done for the *Results* subsection *Effect of membrane composition on lipid block* are listed in *SI Table 2*.

The coarse-grained systems were energy minimized and equilibrated for 500 ns to reach the temperature of 323 K and pressure of 1 bar; for equilibration, v-rescale thermostat ^80^ and Berendsen barostat ^84^. Martini3 parameters with elastic networks between backbone beads were used (500 kJ*mol^-1^*nm^-2^, with the lower and upper elastic bond cut-off lengths 0.5 and 0.9 nm, respectively, including intersubunit elastic bonds) ^49,50^. During equilibration, restraints with a force constant of 1000 kJ*mol^-1^*nm^-2^ on the backbone beads of the protein were used; for production runs it was decreased to 1 kJ*mol^-1^*nm^-2^. Lennard-Jones interactions were treated with a cutoff such that the potential was 0 at 1.1 nm. Electrostatic interactions were treated using reaction field with a 1.1 nm cut-off, with a dielectric constant of 15 before the cut-off, and infinity beyond it. V-rescale thermostat and Parrinello-Rahman barostat were used for production runs. The simulation timestep in Martini3 simulations was 20 fs. For each combination of bilayer composition and channel state, a 40 µs-long simulation was run. In order to prevent water beads from entering the pore in the respective simulations, a spherical flat-bottomed repulsive position restraint with a radius of 1.1 nm, centered on the center of mass of I301 backbone beads was used; initially, the pore contained no water beads.

### Data analysis

Pore hydration, lipid entry into the pore, lipid interaction with the protein residues and conformational changes were analyzed using Python3 and mdanalysis ^85,86^. For the pore occupancy analysis, the pore was defined as a cylinder with a 10 Å radius in the membrane plane, and 20 Å depth relative to the center of mass of T276 which form the lower S4 K^+^ binding site in the SF. To count permeation events and calculate conductances, a custom Fortran script was used ^87^. An outward permeation event was defined as a transition of a K^+^ from the S2 to the S3 binding site in the SF (S3 to S2 for inward permeations). To calculate conductances, the difference between the number of outward and inward permeation events was used. Errors were calculated as 95% confidence intervals using Student’s *t-*distribution. Electrostatic potential maps were obtained using the g_elpot tool ^88^. The data were plotted using matplotlib and seaborn ^89,90^. Molecular visualizations were rendered using VMD ^91^.

## Acknowledgements and funding sources

The authors acknowledge funding from the Deutsche Forschungsgemeinschaft (DFG, German Research Foundation) through FOR2518 ‘Dynion’ Project P5.

## Author contributions

B.L.d.G. and W.K. designed and supervised the project. A.M. performed molecular dynamics simulations and analyzed the data. A.M, B.L.d.G. and W.K. wrote the manuscript.

---

**SI Fig. 1.**
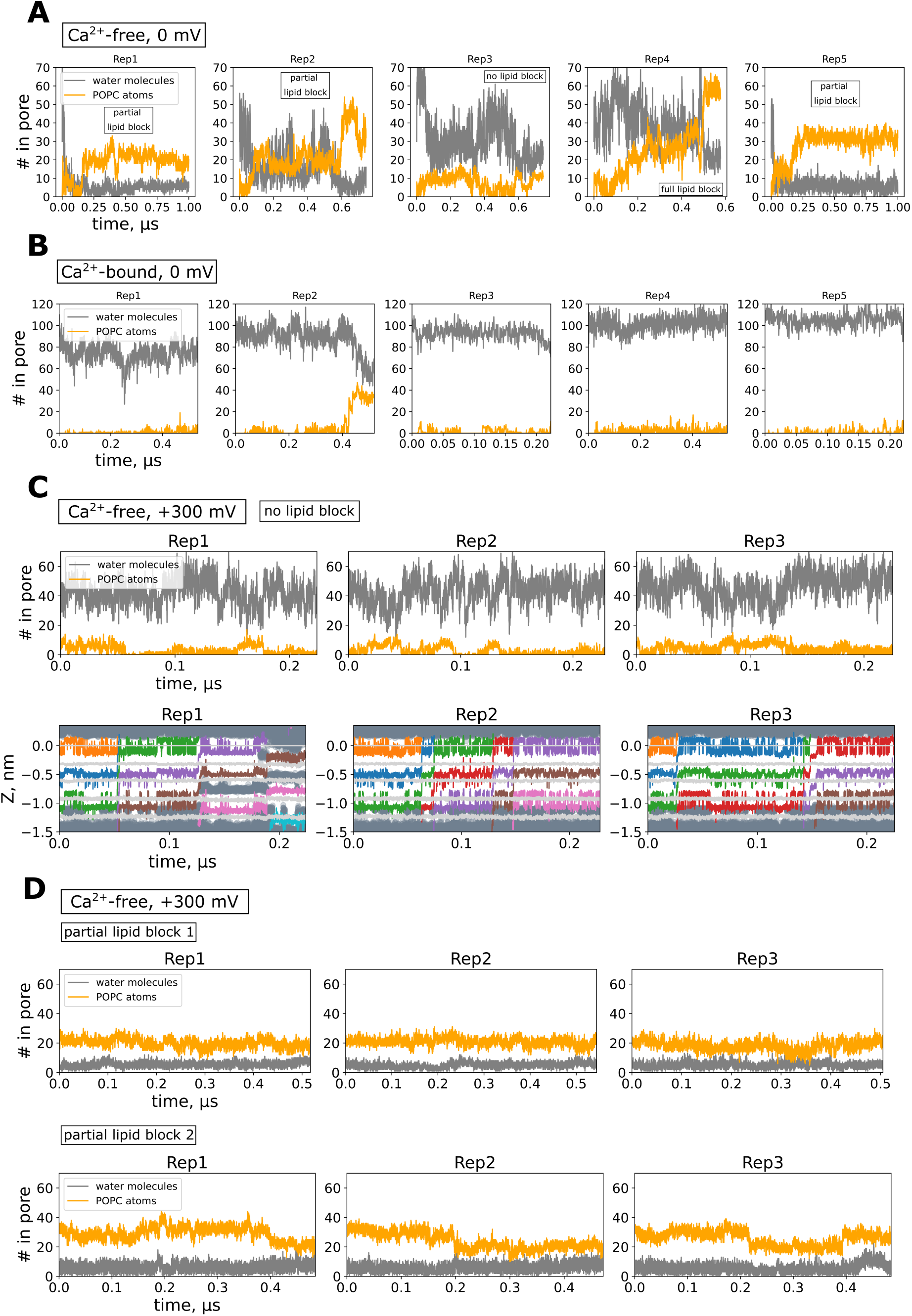
Time traces of the number of lipid atoms (non-hydrogen atoms only; each lipid contains ∼50 non-hydrogen atoms) and water molecules in the pore in atomistic simulations without applied voltage of (*A*) Ca^2+^-free, where the pore was mostly dehydrated and blocked by lipids, except for one replica (‘Rep4’) which showed no lipid block and an overall higher level of hydration; (*B*) Ca^2+^-bound, where the pore was mostly hydrated, except for one replica with some decrease in pore hydration (‘Rep4’), and (*C, upper*) replicas of Ca^2+^-free under applied voltage, where the pore remained hydrated, not blocked by lipids, and conductive, as indicated by time traces of K^+^ (colored lines), water (dark gray scatter) in the SF (carbonyls and hydroxyls of SF-forming K^+^ binding sites are shown as light gray lines) (*C, lower*). (*D*) Time traces of lipid and water oxygen atoms in the pore, Ca^2+^-free +300 mV simulation sets with partial lipid block.

**SI Fig. 2.**
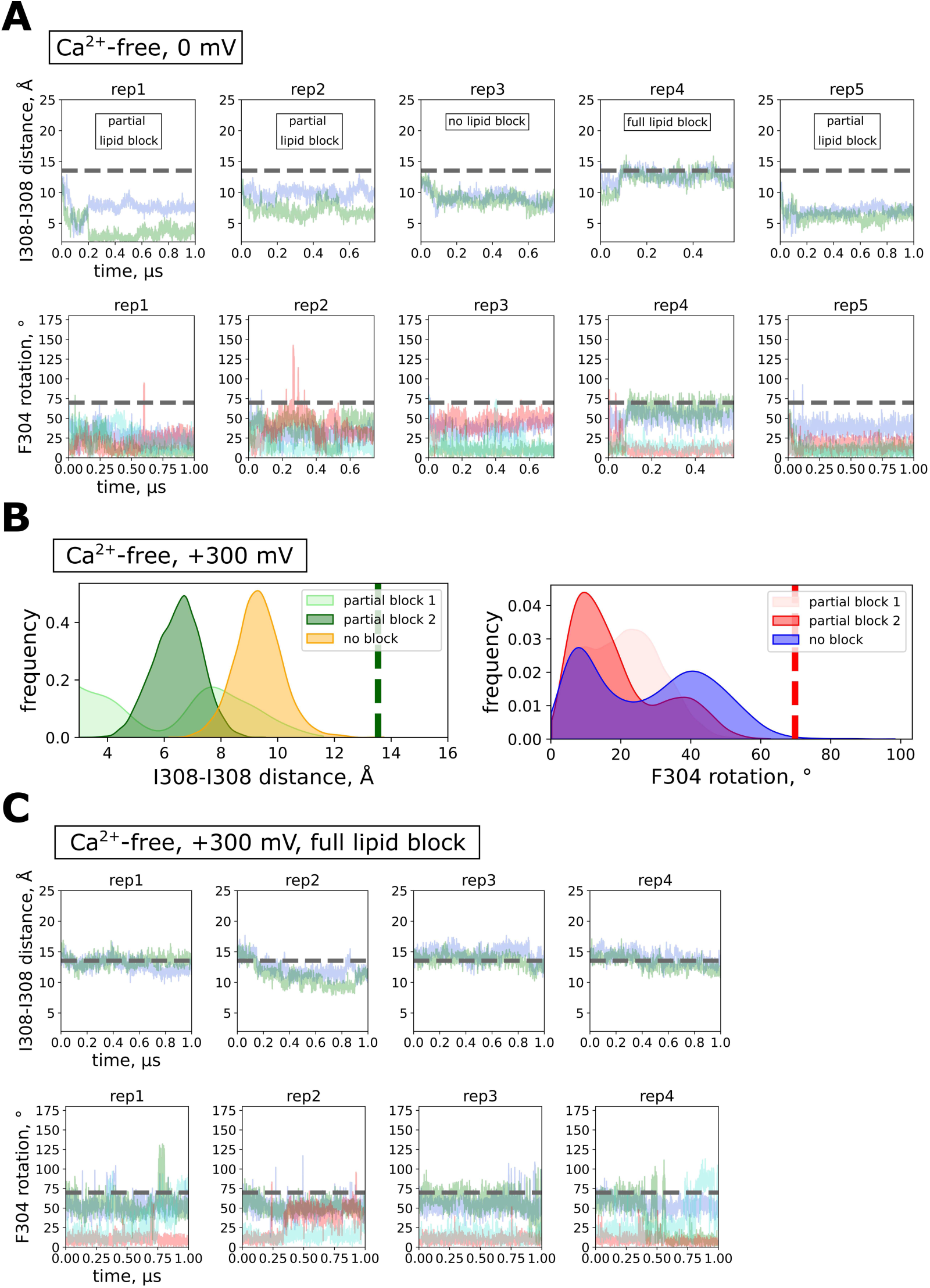
(*A*) Time traces of pore opening (distance between CA atoms of I308 residues of opposing subunits) and inward rotation of F304 (defined as the angle between center of mass of all F304 and CA, CZ atoms of F304 for each subunit) in the Ca^2+^-free state, in atomistic simulations without applied voltage. (*B*) Pore closure and F304 rotation in +300 mV simulations of Ca^2+^-free in the two simulation sets with a partial lipid block, and the simulation set where the lipid block did not occur. (*C*) Time traces of pore opening and inward rotation of F304 in the Ca^2+^-free state, in atomistic simulations at +300 mV where full lipid block occurred. Dashed lines represent the CryoEM structure values for the Ca^2+^-free state.

**SI Fig. 3.**
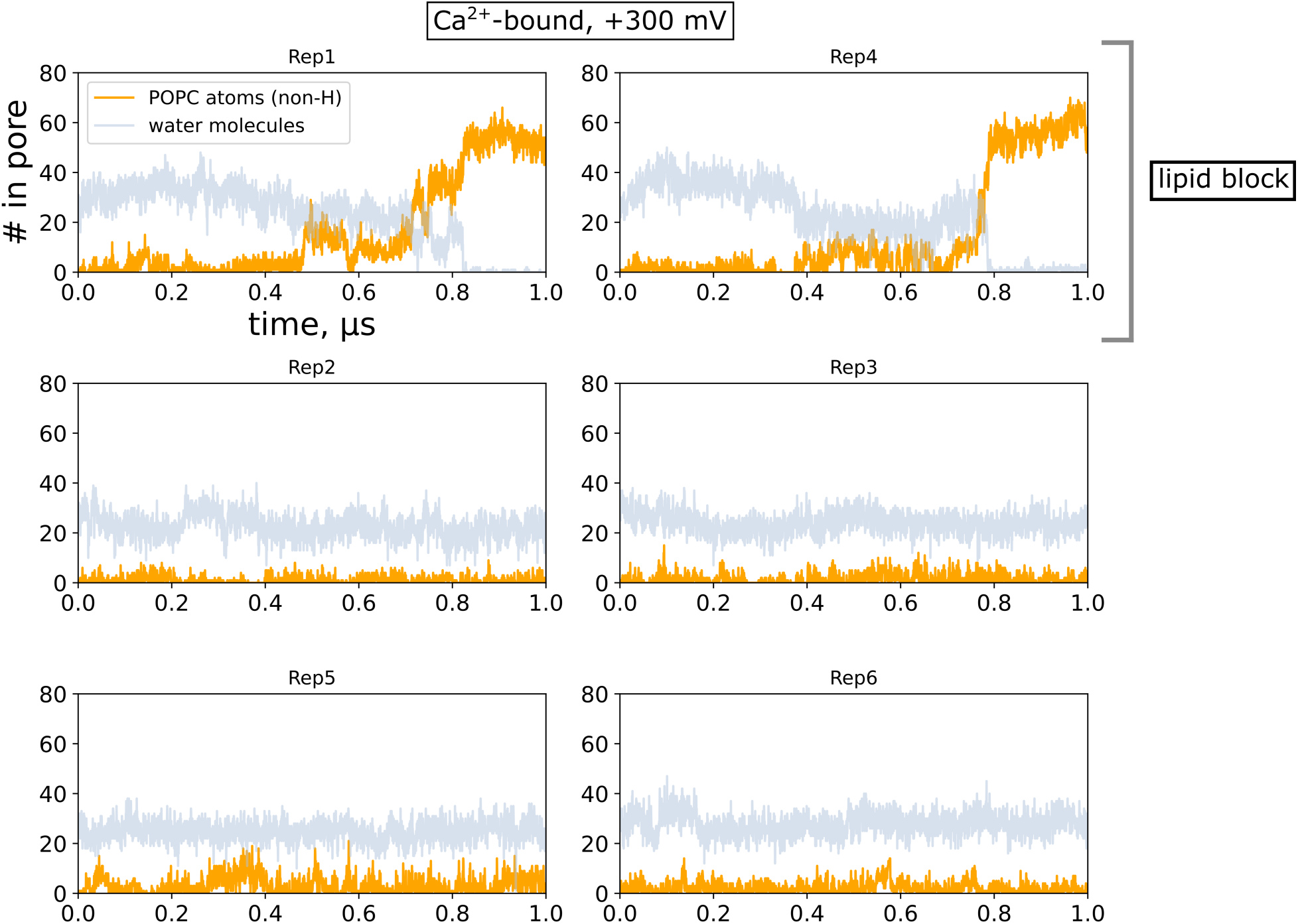
Traces of the number of water molecules and POPC atoms (non-hydrogen, out of ∼50 per lipid molecule) in the pore in the atomistic simulations of the Ca^2+^-bound state at +300 mV. The replicas where the lipid block of the pore occurred are highlighted.

**SI Fig. 4.**
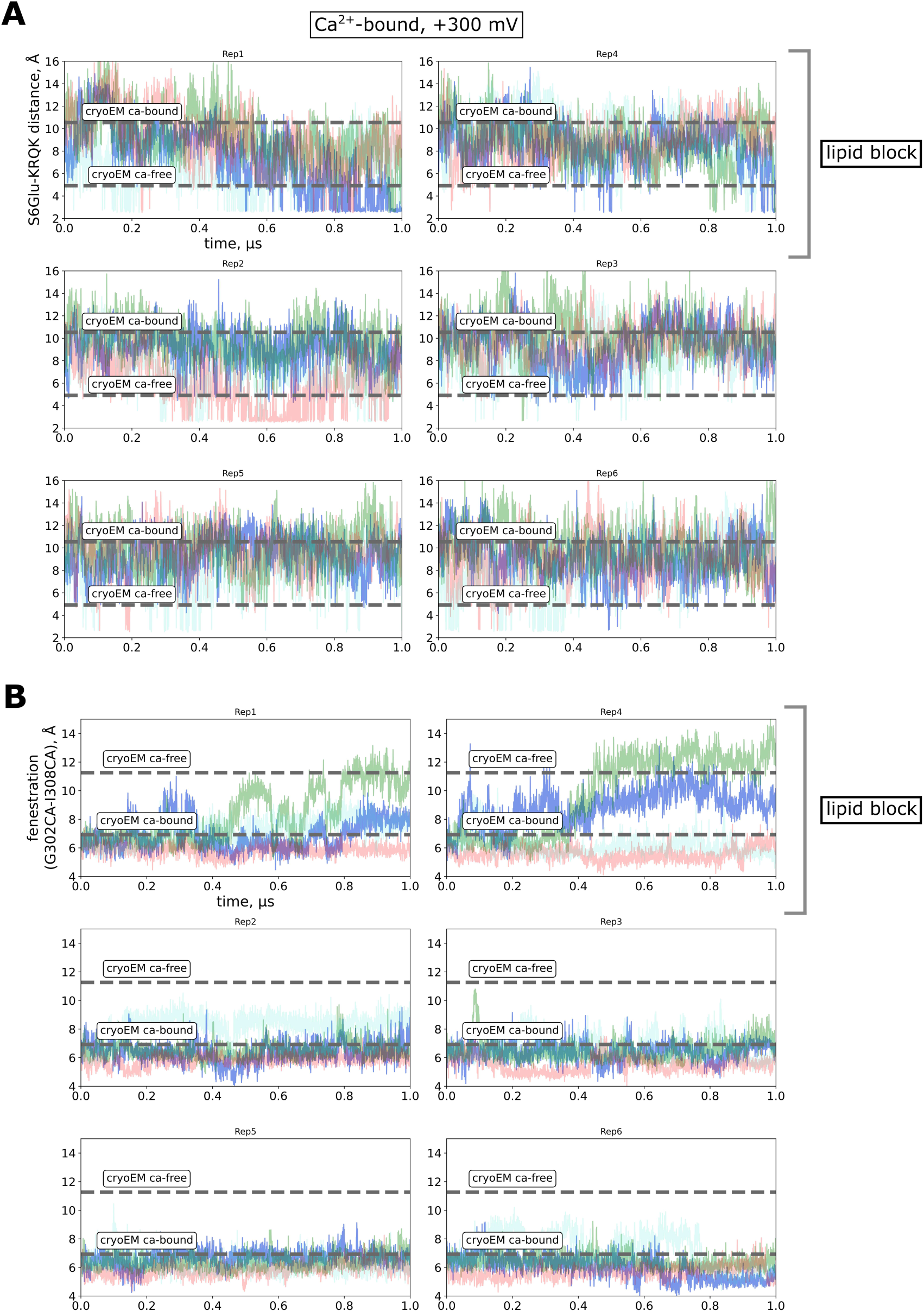
Atomistic simulations of the Ca^2+^-bound state at +300 mV. Time traces of (*A*) distances between E310/E313 (‘S6Glu’) and ^317^KRQK^320^ of neighboring subunits, illustrating the salt bridge formation between the two charged clusters at the end of S6 helices, and (*B*) distances between CA atoms of G302 and I308 of neighboring subunits, illustrating the widening of fenestrations between S6 helices; different colors represent distances between different residue pairs (4 in total). Simulation replicas where spontaneous lipid block occurred (‘Rep1’ and ‘Rep4’) are indicated.

**SI Fig. 5.**
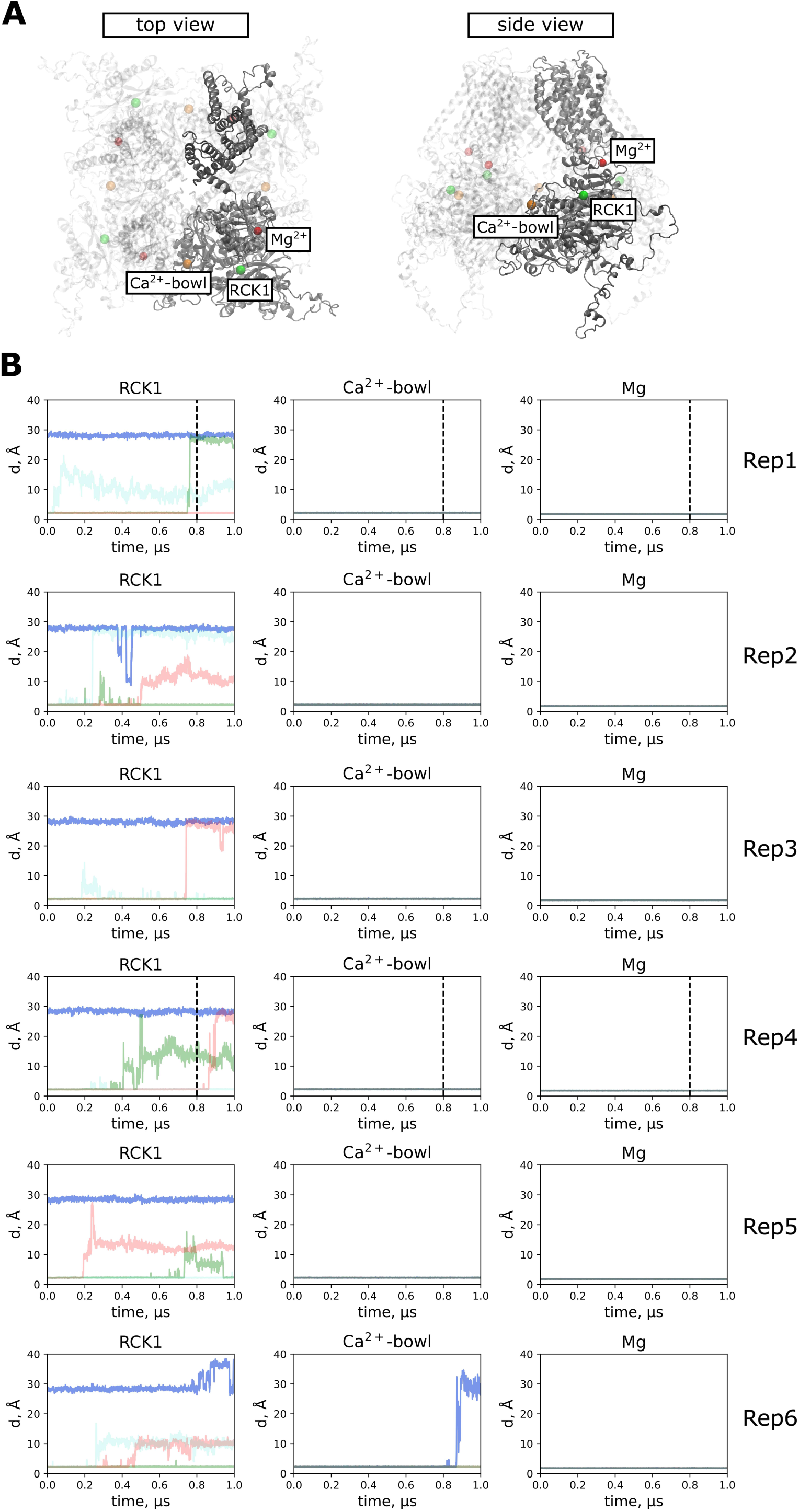
(*A*) BK contains two Ca^2+^-binding sites - RCK1 (green) and Ca^2+^-bowl (orange), as well as one Mg^2+^-binding site (red) per subunit. The ion binding sites belonging to one of the four subunits are shown as non-transparent, and transparent for other subunits. (*B*) Time traces of the distance from a given binding site to the nearest Ca^2+^ or Mg^2+^ respectively, illustrating spontaneous unbinding of Ca^2+^ (each color corresponds to one subunit), simulations of Ca^2+^-bound under +300 mV. Dashed vertical lines in replicas Rep1 and Rep4 indicate time when full lipid block happened

**SI Fig. 6.**
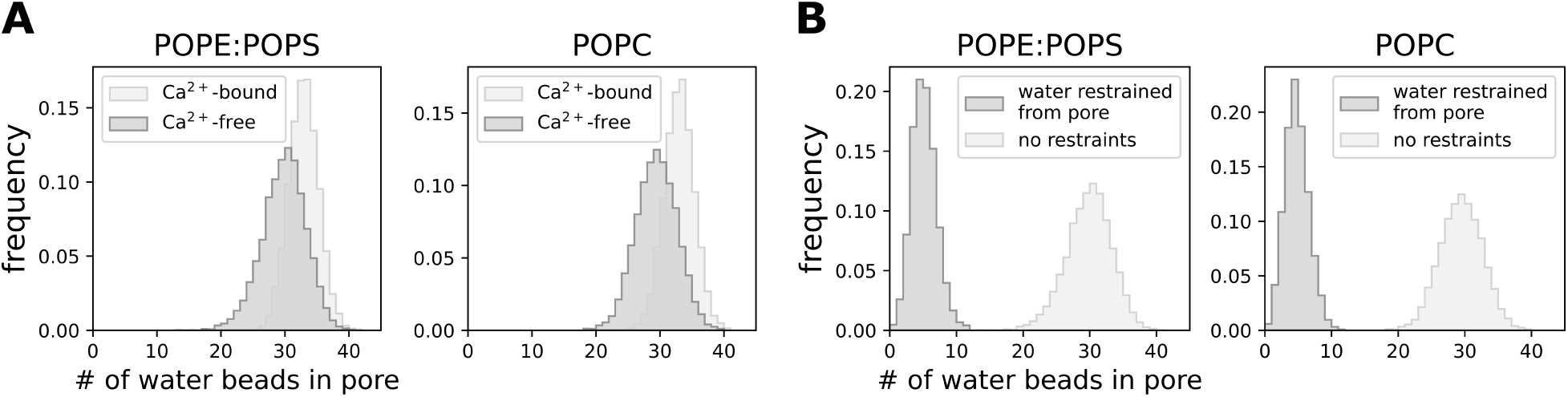
Distribution of water beads in the pore (*A*) without, and (*B*) with restraints on water preventing it from entering the pore in Martini3 simulations.

**SI Fig. 7.**
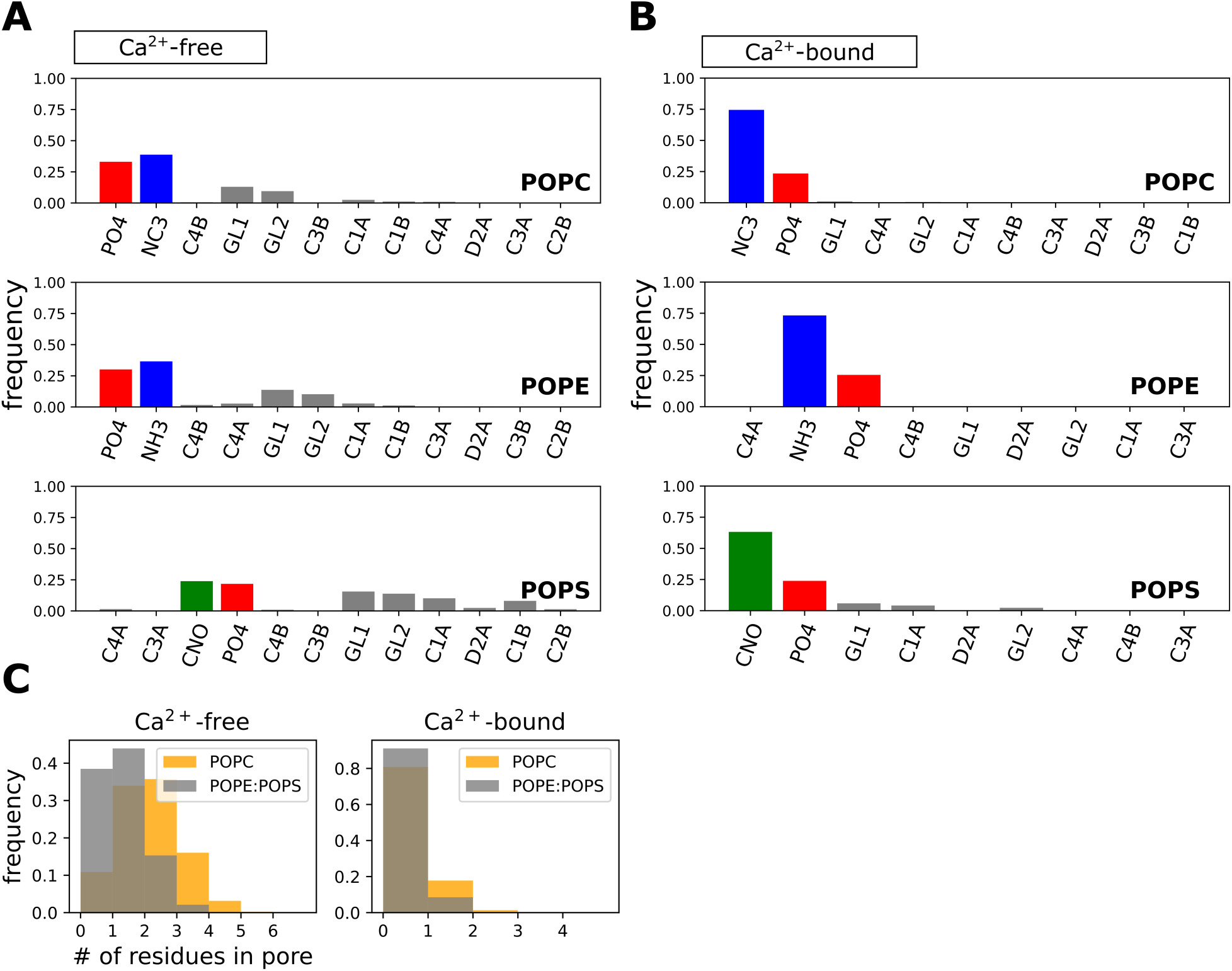
Relative frequencies of lipid beads occupying the pore in (*A*) Ca^2+^-free (hydrated pore) and (*B*) Ca^2+^-bound states in Martini3 simulations; colored bars represent the headgroup beads (excluding the glycerol beads), blue corresponds to positively charged beads, red to negative and green to non-charged. (*C*) Distribution of the number of individual lipid residues simultaneously occupying the pore in Martini3 simulations.

**SI Fig. 8.**
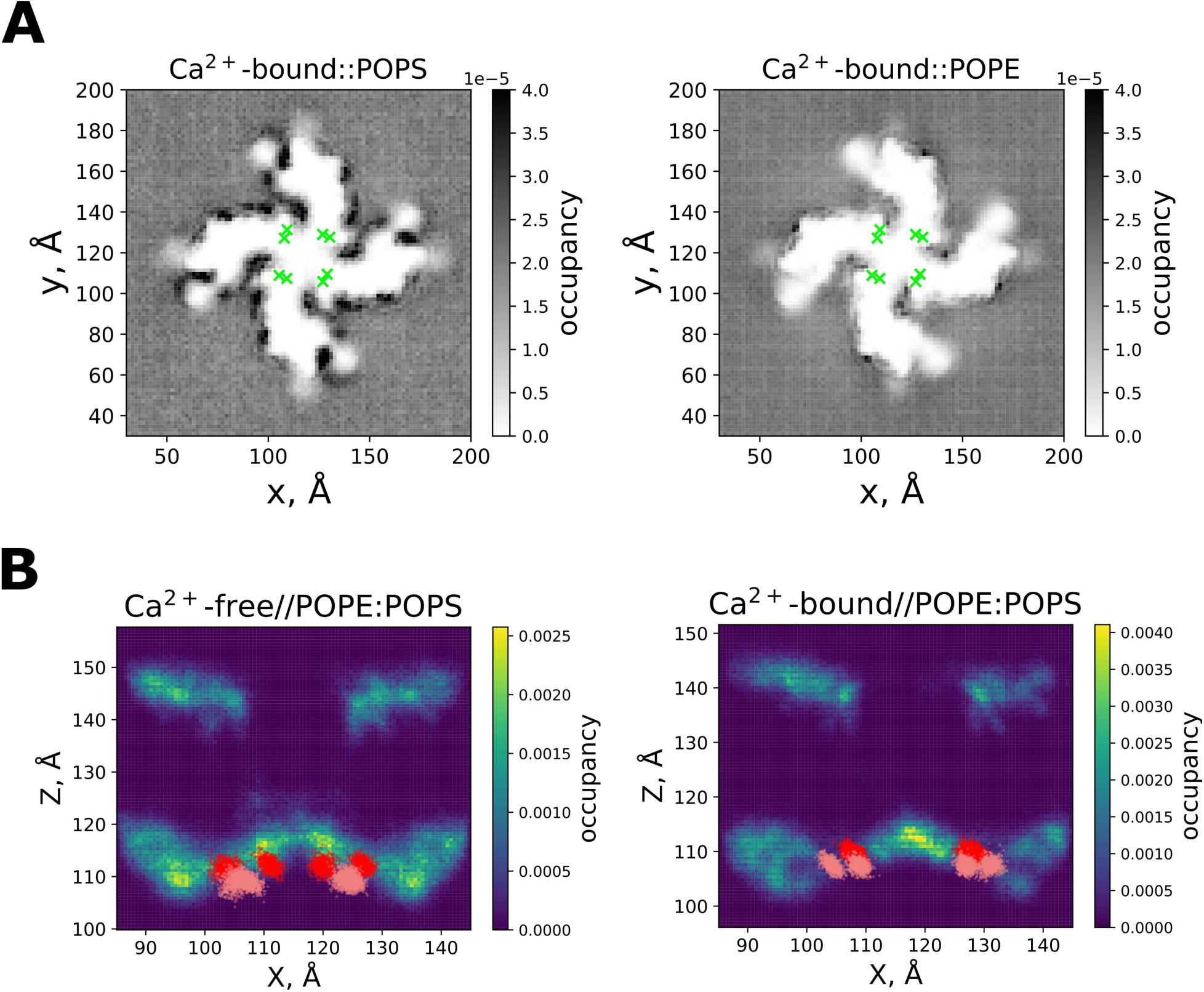
(*A*) Lipid distribution around the channel in Martini3 simulations (Ca^2+^-bound state); E310/E313 residues in S6 helices are shown as reference (green markers). (*B*) Occupancies of PO4 beads (corresponding to phosphate groups; only beads within 3 nm of E310/E313 in the membrane plane were counted), shown in a plane perpendicular to the membrane. Note that the densities are integrated along the Y axis, and the density in the pore region of the channel (X = 110..120 Å) does not necessarily represent PO4 beads occupying the pore.

**SI Fig. 9.**
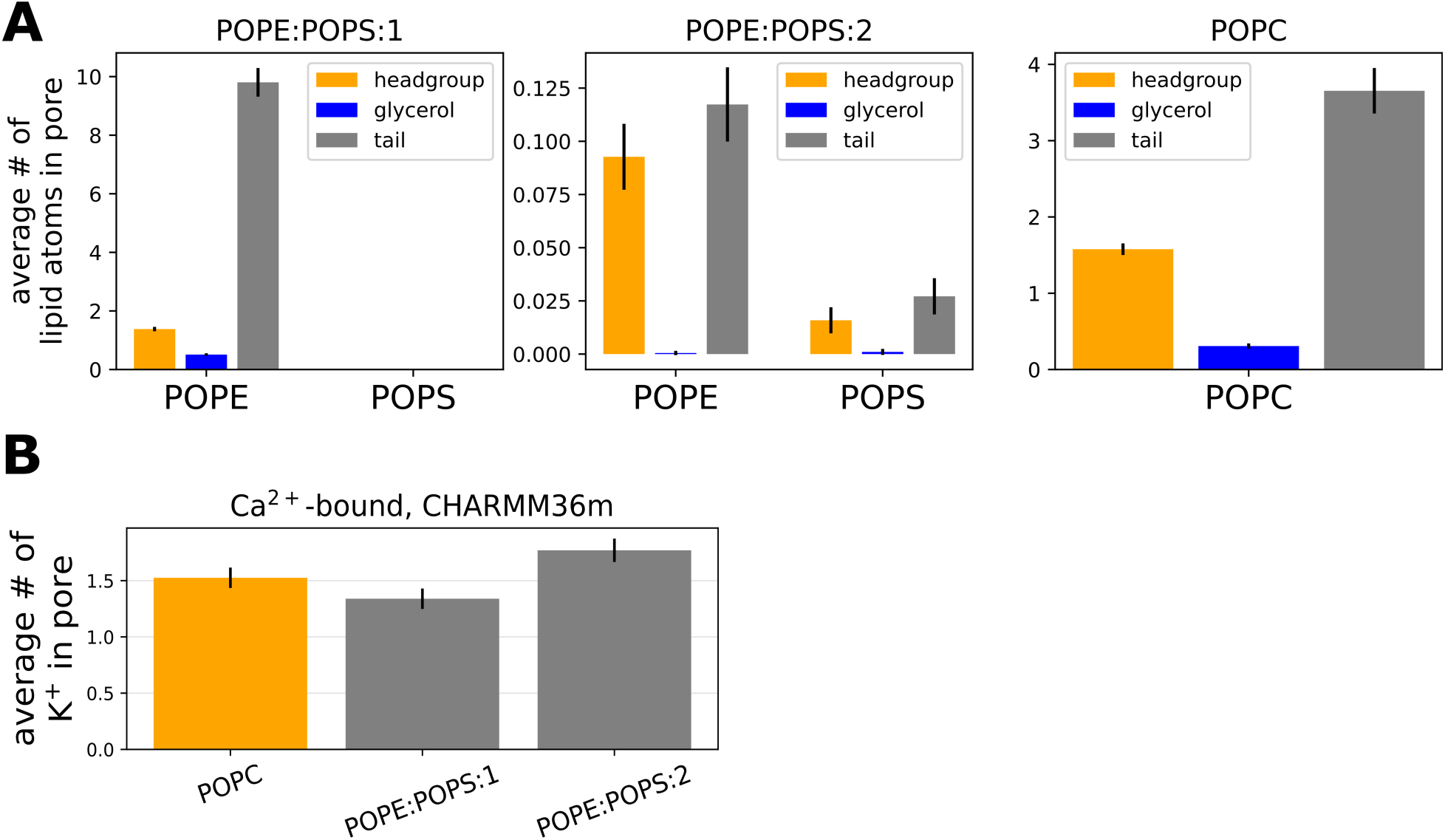
(*A*) Occupancy of the pore by atoms belonging to either the headgroup (excluding glycerol), glycerol or the tail of a given lipid, and (*B*) average number of K^+^ ions in the pore in atomistic simulations (Ca^2+^-bound state, 0 mV).

**SI Fig. 10.**
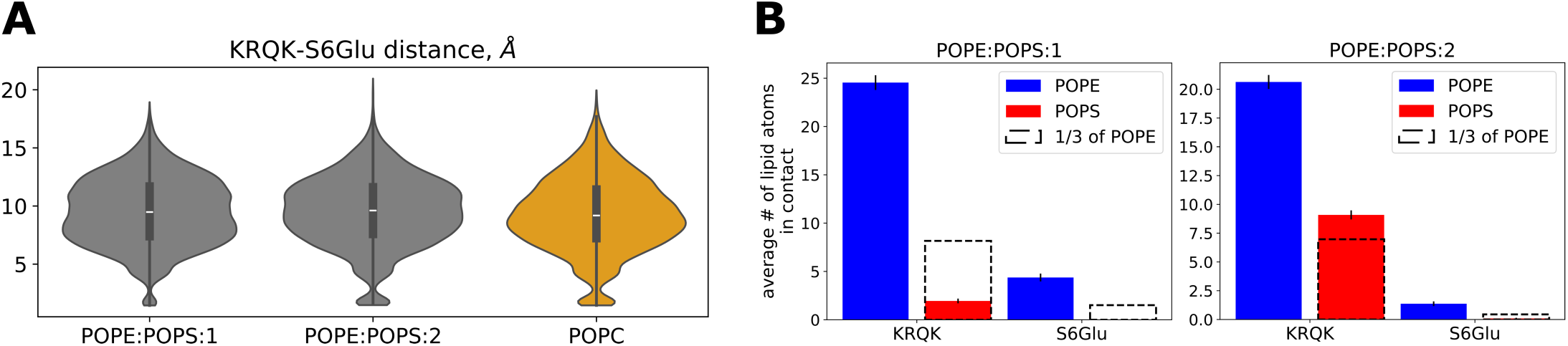
Atomistic simulations of the full-length channel (Ca^2+^-bound state) in POPC compared to POPE:POPS membranes. (*A*) Distributions of distances between ^317^KRQK^320^ and E310/E313 (‘S6Glu’) residues of neighboring subunits. (*B*) Average number of contacts between headgroups belonging to a given lipid type, and ^317^KRQK^320^ or S6Glu in atomistic simulations of Ca^2+^-bound in POPE:POPS. POPE:POPS:1 and POPE:POPS:2 indicate simulations started from the two different initial configurations of the membrane.

**SI Fig. 11.**
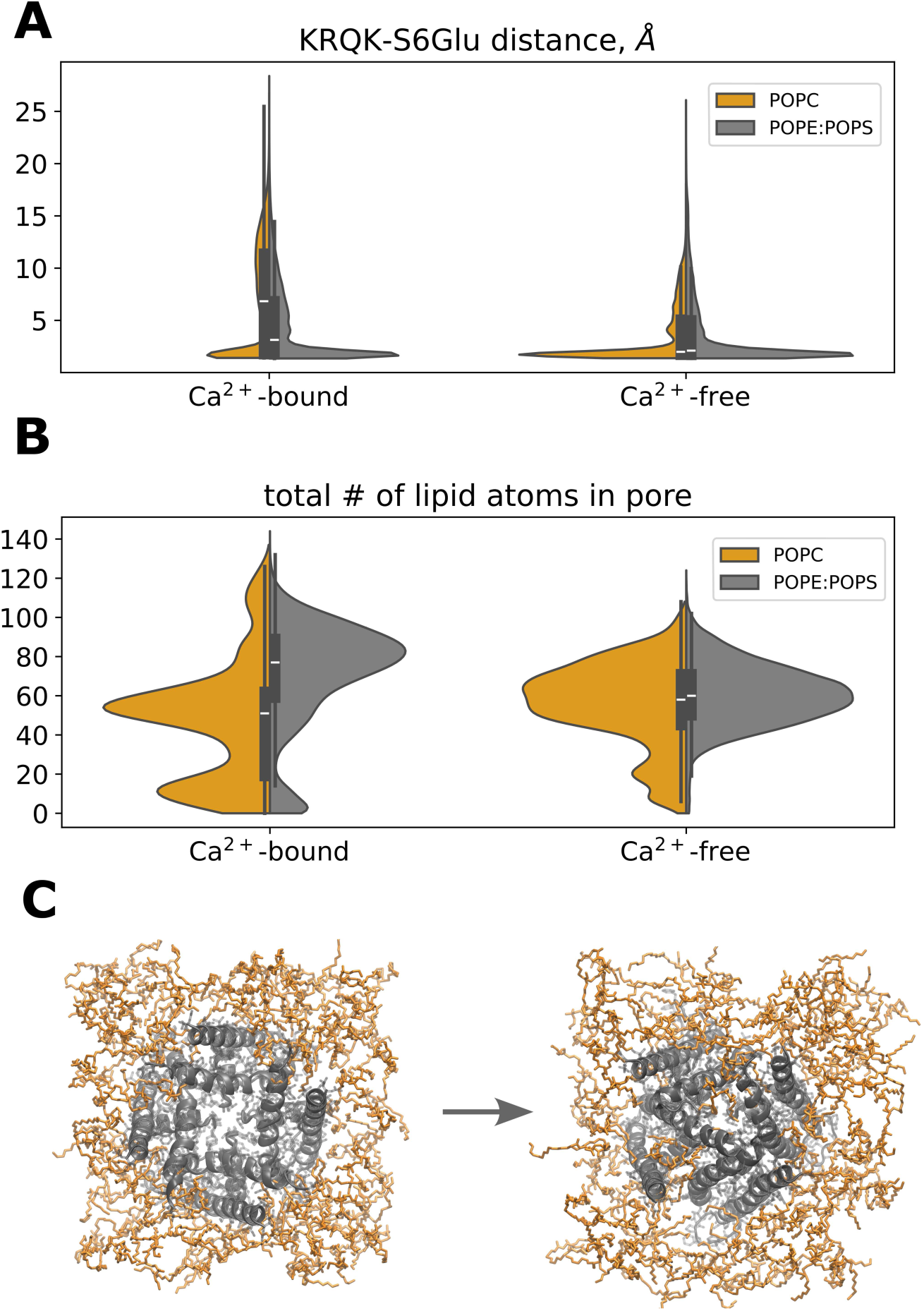
Atomistic simulations of the isolated pore domain. Distributions of (*A*) distances between ^317^KRQK^320^ and E310/E313, and (*B*) number of lipid atoms (non-hydrogen, out of ∼50 per lipid molecule) in the pore. (*C*) Simulation snapshots showing low structural stability of the isolated pore domain and high propensity to be blocked by lipids regardless of the state of the channel (here, Ca^2+^-bound is shown).

**SI Fig. 12.**
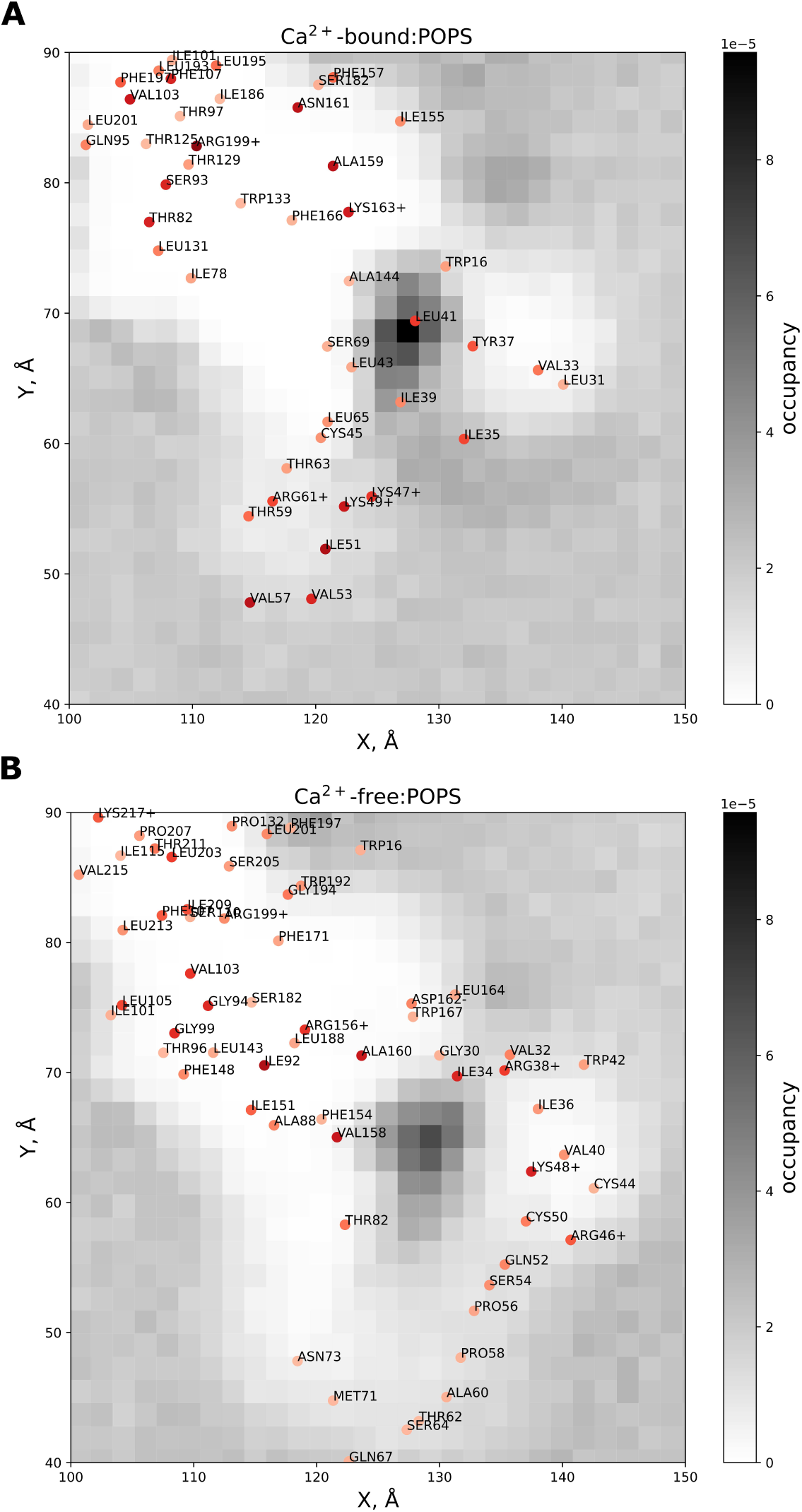
Occupancies of PO4 beads of POPS near the binding site located in the voltage sensor in (*A*) Ca^2+^-bound and (*B*) Ca^2+^-free states, with protein residues interacting with POPS highlighted; darker color indicates more frequent contacts.

**SI Table 1.** List of simulated systems discussed in the ‘Results’ subsections ‘All-atom simulations of Ca^2+^-bound and Ca^2+^-free states’ and ‘Spontaneous closure of the open state’.

| Channel structure | Membrane composition | Force field | Voltage | Duration |
| --- | --- | --- | --- | --- |
| Ca<sup>2+</sup>-bound | POPC | CHARMM36m | 0 mV | 5x0.2..0.5 μs |
| | | | +300 mV | 6x1 μs (started from of one of the 0 mV replicas) |
| Ca<sup>2+</sup>-free | POPC | CHARMM36m | 0 mV | 5x0.2..0.5 μs |
| | | | +300 mV | 3x0.2 μs (started from a 0 mV replica with no lipid block), 6x0.6..0.7 μs (started from two 0 mV replicas with partial lipid block, with 3 applied voltage replicas for each initial frame), 4x1 μs (started from a 0 mV replica with full lipid block) |

**SI Table 2.** List of simulated systems discussed in the ‘Results’ subsection ‘Effect of membrane composition on lipid block’. Unless noted, listed simulations correspond to those of a full-length channel.

|  |  |  |  |  |
| --- | --- | --- | --- | --- |
| Ca<sup>2+</sup>-bound | POPE:POPS,<br>3:1 | Martini3 | 0 mV | 40 μs |
| Ca<sup>2+</sup>-free | POPE:POPS,<br>3:1 | Martini3 | 0 mV | 40 μs (no restraints on water), ~20 μs (with restraints on water molecules preventing them from entering the pore) |
| Ca<sup>2+</sup>-bound | POPC | Martini3 | 0 mV | 40 μs |
| Ca<sup>2+</sup>-free | POPC | Martini3 | 0 mV | 40 μs (no restraints on water), ~20 μs (with restraints on water molecules preventing them from entering the pore) |
| Ca<sup>2+</sup>-bound | POPE:POPS,<br>3:1 | CHARMM36m | 0 mV | 4x0.2..0.5 μs (initial membrane configuration 1),<br>4x0.3..0.4 μs (initial membrane configuration 2) |
| Ca<sup>2+</sup>-free | POPE:POPS,<br>3:1 | CHARMM36m | 0 mV | 4x0.2..0.5 μs (initial membrane configuration 1),<br>4x0.2..0.3 μs (initial membrane configuration 2) |
| Ca<sup>2+</sup>-bound,<br>isolated pore domain | POPC | CHARMM36m | 0 mV | 10x1 μs |
| Ca<sup>2+</sup>-free,<br>isolated pore domain | POPC | CHARMM36m | 0 mV | 10x1 μs |
| Ca<sup>2+</sup>-bound,<br>isolated pore<br>domain | POPE:POPS,<br>3:1 | CHARMM36m | 0 mV | 4 different initial<br>membrane<br>configurations,<br>5x1 μs each |
| Ca<sup>2+</sup>-free,<br>isolated pore<br>domain | POPE:POPS,<br>3:1 | CHARMM36m | 0 mV | 4 different initial<br>membrane<br>configurations,<br>5x1 μs each |

## References

1. González, C. et al. K(+) channels: function-structural overview. Compr. Physiol. 2, 2087–2149 (2012).

2. Hibino, H. et al. Inwardly rectifying potassium channels: their structure, function, and physiological roles. Physiol. Rev. 90, 291–366 (2010).

3. Jespersen, T., Grunnet, M. & Olesen, S.-P. The KCNQ1 potassium channel: from gene to physiological function. Physiology (Bethesda) 20, 408–416 (2005).

4. Latorre, R., Oberhauser, A., Labarca, P. & Alvarez, O. Varieties of calcium-activated potassium channels. Annu. Rev. Physiol. 51, 385–399 (1989).

5. Latorre, R. et al. Molecular Determinants of BK Channel Functional Diversity and Functioning. Physiol. Rev. 97, 39–87 (2017).

6. Contreras, G. F. et al. A BK (Slo1) channel journey from molecule to physiology. Channels (Austin) 7, 442–458 (2013).

7. González-Sanabria, N., Echeverría, F., Segura, I., Alvarado-Sánchez, R. & Latorre, R. BK in double-membrane organelles: A biophysical, pharmacological, and functional survey. Front. Physiol. 12, 761474 (2021).

8. Hite, R. K., Tao, X. & MacKinnon, R. Structural basis for gating the high-conductance Ca2+-activated K+ channel. Nature 541, 52–57 (2017).

9. Tao, X., Hite, R. K. & MacKinnon, R. Cryo-EM structure of the open high-conductance Ca2+-activated K+ channel. Nature 541, 46–51 (2017).

10. Tao, X., Zhao, C. & MacKinnon, R. Membrane protein isolation and structure determination in cell-derived membrane vesicles. Proc. Natl. Acad. Sci. U. S. A. 120, e2302325120 (2023).

11. Tao, X. & MacKinnon, R. Molecular structures of the human Slo1 K+ channel in complex with β4. Elife 8, (2019).

12. Contreras, G. F., Sheng, R., Latorre, R. & Perozo, E. Structural basis of voltage-dependent gating in BK channels and its coupling to the calcium sensor. bioRxiv (2023) doi:10.1101/2023.12.29.573674.

13. del Camino, D. & Yellen, G. Tight steric closure at the intracellular activation gate of a voltage-gated K+ channel. Neuron 32, 649–656 (2001).

14. Labro, A. J. & Snyders, D. J. Being flexible: the voltage-controllable activation gate of kv channels. Front. Pharmacol. 3, 168 (2012).

15. Moldenhauer, H., Díaz-Franulic, I., González-Nilo, F. & Naranjo, D. Effective pore size and radius of capture for K(+) ions in K-channels. Sci. Rep. 6, 19893 (2016).

16. Mironenko, A., Zachariae, U., de Groot, B. L. & Kopec, W. The Persistent Question of Potassium Channel Permeation Mechanisms. J. Mol. Biol. 433, 167002 (2021).

17. Lin, H., Li, J., Zhang, Q., Yang, H. & Chen, S. C-type inactivation and proton modulation mechanisms of the TASK3 channel. Proc. Natl. Acad. Sci. U. S. A. 121, e2320345121 (2024).

18. Schewe, M. et al. A non-canonical voltage-sensing mechanism controls gating in K2P K(+) channels. Cell 164, 937–949 (2016).

19. Tang, Q.-Y., Zeng, X.-H. & Lingle, C. J. Closed-channel block of BK potassium channels by bbTBA requires partial activation. J. Gen. Physiol. 134, 409–436 (2009).

20. Li, W. & Aldrich, R. W. Unique inner pore properties of BK channels revealed by quaternary ammonium block. J. Gen. Physiol. 124, 43–57 (2004).

21. Wilkens, C. M. & Aldrich, R. W. State-independent block of BK channels by an intracellular quaternary ammonium. J. Gen. Physiol. 128, 347–364 (2006).

22. Geng, Y., Niu, X. & Magleby, K. L. Low resistance, large dimension entrance to the inner cavity of BK channels determined by changing side-chain volume. J. Gen. Physiol. 137, 533–548 (2011).

23. Zhou, Y., Xia, X.-M. & Lingle, C. J. Cysteine scanning and modification reveal major differences between BK channels and Kv channels in the inner pore region. Proc. Natl. Acad. Sci. U. S. A. 108, 12161–12166 (2011).

24. Tian, Y., Heinemann, S. H. & Hoshi, T. Large-conductance Ca2+- and voltage-gated K+ channels form and break interactions with membrane lipids during each gating cycle. Proc. Natl. Acad. Sci. U. S. A. 116, 8591–8596 (2019).

25. Aryal, P., Sansom, M. S. P. & Tucker, S. J. Hydrophobic gating in ion channels. J. Mol. Biol. 427, 121–130 (2015).

26. Beckstein, O. & Sansom, M. S. P. Liquid-vapor oscillations of water in hydrophobic nanopores. Proc. Natl. Acad. Sci. U. S. A. 100, 7063–7068 (2003).

27. Hummer, G., Rasaiah, J. C. & Noworyta, J. P. Water conduction through the hydrophobic channel of a carbon nanotube. Nature 414, 188–190 (2001).

28. Zhu, F. & Hummer, G. Drying transition in the hydrophobic gate of the GLIC channel blocks ion conduction. Biophys. J. 103, 219–227 (2012).

29. Sauguet, L. et al. Crystal structures of a pentameric ligand-gated ion channel provide a mechanism for activation. Proc. Natl. Acad. Sci. U. S. A. 111, 966–971 (2014).

30. Miller, S. et al. Domain organization of the MscS mechanosensitive channel of Escherichia coli. EMBO J. 22, 36–46 (2003).

31. Anishkin, A. & Sukharev, S. Water dynamics and dewetting transitions in the small mechanosensitive channel MscS. Biophys. J. 86, 2883–2895 (2004).

32. Sotomayor, M. & Schulten, K. Molecular dynamics study of gating in the mechanosensitive channel of small conductance MscS. Biophys. J. 87, 3050–3065 (2004).

33. Aryal, P., Abd-Wahab, F., Bucci, G., Sansom, M. S. P. & Tucker, S. J. A hydrophobic barrier deep within the inner pore of the TWIK-1 K2P potassium channel. Nat. Commun. 5, 4377 (2014).

34. Jia, Z., Yazdani, M., Zhang, G., Cui, J. & Chen, J. Hydrophobic gating in BK channels. Nat. Commun. 9, 3408 (2018).

35. Gu, R.-X. & de Groot, B. L. Central cavity dehydration as a gating mechanism of potassium channels. Nat. Commun. 14, 2178 (2023).

36. Nordquist, E. B., Jia, Z. & Chen, J. Inner pore hydration free energy controls the activation of big potassium channels. Biophys. J. 122, 1158–1167 (2023).

37. Chen, X. & Aldrich, R. W. Charge substitution for a deep-pore residue reveals structural dynamics during BK channel gating. J. Gen. Physiol. 138, 137–154 (2011).

38. Chen, X., Yan, J. & Aldrich, R. W. BK channel opening involves side-chain reorientation of multiple deep-pore residues. Proc. Natl. Acad. Sci. U. S. A. 111, E79–88 (2014).

39. Nordquist, E. et al. Incorporating physics to overcome data scarcity in predictive modeling of protein function: A case study of BK channels. PLoS Comput. Biol. 19, e1011460 (2023).

40. Coronel, L., Di Muccio, G., Rothberg, B. S., Giacomello, A. & Carnevale, V. Lipid-mediated hydrophobic gating in the BK potassium channel. Nat. Commun. 16, 7354 (2025).

41. Yuan, C., O’Connell, R. J., Jacob, R. F., Mason, R. P. & Treistman, S. N. Regulation of the gating of BKCa channel by lipid bilayer thickness. J. Biol. Chem. 282, 7276–7286 (2007).

42. Vaithianathan, T. et al. Direct regulation of BK channels by phosphatidylinositol 4,5-bisphosphate as a novel signaling pathway. J. Gen. Physiol. 132, 13–28 (2008).

43. Crowley, J. J., Treistman, S. N. & Dopico, A. M. Distinct structural features of phospholipids differentially determine ethanol sensitivity and basal function of BK channels. Mol. Pharmacol. 68, 4–10 (2005).

44. Hui, C., de Vries, R., Kopec, W. & de Groot, B. Charge Scaling in Potassium Channel Simulations: Conductance, Ion Occupancy, Voltage Response, and Selectivity. Biophysics (2024).

45. Talukder, G. & Aldrich, R. W. Complex voltage-dependent behavior of single unliganded calcium-sensitive potassium channels. Biophys. J. 78, 761–772 (2000).

46. Contreras, G. F., Shen, R., Latorre, R. & Perozo, E. Structural basis of voltage-dependent gating in BK channels. Nat. Commun. 16, 5846 (2025).

47. Dopico, A. M. & Bukiya, A. N. Lipid regulation of BK channel function. Front. Physiol. 5, 312 (2014).

48. Vaithianathan, T., Schneider, E. H., Bukiya, A. N. & Dopico, A. M. Cholesterol and PIP2 modulation of BKCa channels. Adv. Exp. Med. Biol. 1422, 217–243 (2023).

49. Souza, P. C. T. et al. Martini 3: a general purpose force field for coarse-grained molecular dynamics. Nat. Methods 18, 382–388 (2021).

50. Periole, X., Cavalli, M., Marrink, S.-J. & Ceruso, M. A. Combining an elastic network with a coarse-grained molecular force field: Structure, dynamics, and intermolecular recognition. J. Chem. Theory Comput. 5, 2531–2543 (2009).

51. Schewe, M. et al. A pharmacological master key mechanism that unlocks the selectivity filter gate in K+ channels. Science 363, 875–880 (2019).

52. Woltz, R. L. et al. Atomistic mechanisms of the regulation of small-conductance Ca2+-activated K+ channel (SK2) by PIP2. Proc. Natl. Acad. Sci. U. S. A. 121, e2318900121 (2024).

53. Tian, Y. et al. Two distinct effects of PIP2 underlie auxiliary subunit-dependent modulation of Slo1 BK channels. J. Gen. Physiol. 145, 331–343 (2015).

54. Sun, J. & MacKinnon, R. Structural basis of human KCNQ1 modulation and gating. Cell 180, 340–347.e9 (2020).

55. Ma, D. et al. Structural mechanisms for the activation of human cardiac KCNQ1 channel by electro-mechanical coupling enhancers. Proc. Natl. Acad. Sci. U. S. A. 119, e2207067119 (2022).

56. Tan, X.-F. et al. Structure of the Shaker Kv channel and mechanism of slow C-type inactivation. Sci. Adv. 8, eabm7814 (2022).

57. Cuello, L. G., Jogini, V., Cortes, D. M. & Perozo, E. Structural mechanism of C-type inactivation in K(+) channels. Nature 466, 203–208 (2010).

58. Lippiat, J. D., Standen, N. B. & Davies, N. W. A residue in the intracellular vestibule of the pore is critical for gating and permeation in Ca2+-activated K+ (BKCa) channels. J. Physiol. 529 Pt 1, 131–138 (2000).

59. Carrasquel-Ursulaez, W. et al. Hydrophobic interaction between contiguous residues in the S6 transmembrane segment acts as a stimuli integration node in the BK channel. J. Gen. Physiol. 145, 61–74 (2015).

60. Chang, H. M., Reitstetter, R. & Gruener, R. Lipid-ion channel interactions: increasing phospholipid headgroup size but not ordering acyl chains alters reconstituted channel behavior. J. Membr. Biol. 145, 13–19 (1995).

61. Schmidpeter, P. A. M. et al. Membrane phospholipids control gating of the mechanosensitive potassium leak channel TREK1. Nat. Commun. 14, 1077 (2023).

62. Ma, Q. et al. Insights into the structure and modulation of human TWIK-2. Nat. Commun. 1–14 (2026).

63. Jorgensen, C. et al. Lateral fenestrations in K(+)-channels explored using molecular dynamics simulations. Mol. Pharm. 13, 2263–2273 (2016).

64. McManus, O. B. & Magleby, K. L. Kinetic states and modes of single large-conductance calcium-activated potassium channels in cultured rat skeletal muscle. J. Physiol. 402, 79–120 (1988).

65. Fan, C. et al. Ball-and-chain inactivation in a calcium-gated potassium channel. Nature 580, 288–293 (2020).

66. Paulo, G. et al. Hydrophobically gated memristive nanopores for neuromorphic applications. Nat. Commun. 14, 8390 (2023).

67. Jensen, M. Ø. et al. Principles of conduction and hydrophobic gating in K+ channels. Proc. Natl. Acad. Sci. U. S. A. 107, 5833–5838 (2010).

68. Clarke, A. L., Petrou, S., Walsh, J. V., Jr & Singer, J. J. Modulation of BK(Ca) channel activity by fatty acids: structural requirements and mechanism of action. Am. J. Physiol. Cell Physiol. 283, C1441–53 (2002).

69. Park, J. B., Kim, H. J., Ryu, P. D. & Moczydlowski, E. Effect of phosphatidylserine on unitary conductance and Ba2+ block of the BK Ca2+-activated K+ channel: re-examination of the surface charge hypothesis: Re-examination of the surface charge hypothesis. J. Gen. Physiol. 121, 375–397 (2003).

70. Jo, S., Kim, T., Iyer, V. G. & Im, W. CHARMM-GUI: a web-based graphical user interface for CHARMM. J. Comput. Chem. 29, 1859–1865 (2008).

71. Lee, J. et al. CHARMM-GUI input generator for NAMD, GROMACS, AMBER, OpenMM, and CHARMM/OpenMM simulations using the CHARMM36 additive force field. J. Chem. Theory Comput. 12, 405–413 (2016).

72. Ostmeyer, J., Chakrapani, S., Pan, A. C., Perozo, E. & Roux, B. Recovery from slow inactivation in K+ channels is controlled by water molecules. Nature 501, 121–124 (2013).

73. Weingarth, M. et al. Quantitative analysis of the water occupancy around the selectivity filter of a K+ channel in different gating modes. J. Am. Chem. Soc. 136, 2000–2007 (2014).

74. Long, S. B., Tao, X., Campbell, E. B. & MacKinnon, R. Atomic structure of a voltage-dependent K+ channel in a lipid membrane-like environment. Nature 450, 376–382 (2007).

75. Ye, S., Li, Y. & Jiang, Y. Novel insights into K+ selectivity from high-resolution structures of an open K+ channel pore. Nat. Struct. Mol. Biol. 17, 1019–1023 (2010).

76. Kroon, P. C. et al. Martinize2 and Vermouth: Unified framework for topology generation. *arXiv [q-bio.QM]* (2023) doi:10.7554/elife.90627.1.

77. Insane INSert membrANE - A Simple, Versatile Tool for Building Coarse-Grained Simulation Systems. (Github).

78. Abraham, M. J. et al. GROMACS: High performance molecular simulations through multi-level parallelism from laptops to supercomputers. SoftwareX 1-2, 19–25 (2015).

79. Parrinello, M. & Rahman, A. Polymorphic transitions in single crystals: A new molecular dynamics method. J. Appl. Phys. 52, 7182–7190 (1981).

80. Bussi, G., Donadio, D. & Parrinello, M. Canonical sampling through velocity rescaling. J. Chem. Phys. 126, 014101 (2007).

81. Hess, B., Bekker, H., Berendsen, H. J. C. & Fraaije, J. G. E. M. LINCS: A linear constraint solver for molecular simulations. J. Comput. Chem. 18, 1463–1472 (1997).

82. Darden, T., York, D. & Pedersen, L. Particle mesh Ewald: An N⋅log(N) method for Ewald sums in large systems. J. Chem. Phys. 98, 10089–10092 (1993).

83. Gumbart, J., Khalili-Araghi, F., Sotomayor, M. & Roux, B. Constant electric field simulations of the membrane potential illustrated with simple systems. Biochim. Biophys. Acta 1818, 294–302 (2012).

84. Berendsen, H. J. C., Postma, J. P. M., van Gunsteren, W. F., DiNola, A. & Haak, J. R. Molecular dynamics with coupling to an external bath. J. Chem. Phys. 81, 3684–3690 (1984).

85. Gowers, R. et al. MDAnalysis: A python package for the rapid analysis of molecular dynamics simulations. in Proceedings of the Python in Science Conference 98–105 (SciPy, 2016).

86. Michaud-Agrawal, N., Denning, E. J., Woolf, T. B. & Beckstein, O. MDAnalysis: a toolkit for the analysis of molecular dynamics simulations. J. Comput. Chem. 32, 2319–2327 (2011).

87. Kopec, W., Rothberg, B. S. & de Groot, B. L. Molecular mechanism of a potassium channel gating through activation gate-selectivity filter coupling. Nat. Commun. 10, 5366 (2019).

88. Kostritskii, A. Y., Alleva, C., Cönen, S. & Machtens, J.-P. G_elpot: A tool for quantifying biomolecular electrostatics from molecular dynamics trajectories. J. Chem. Theory Comput. 17, 3157–3167 (2021).

89. Hunter, J. D. Matplotlib: A 2D Graphics Environment. Comput. Sci. Eng. 9, 90–95 (2007).

90. Waskom, M. seaborn: statistical data visualization. J. Open Source Softw. 6, 3021 (2021).

91. Humphrey, W., Dalke, A. & Schulten, K. VMD: visual molecular dynamics. J. Mol. Graph. 14, 33–8, 27–8 (1996).

